# NNV024, a novel humanized anti-CD37 antibody with enhanced ADCC and prolonged plasma half-life in human FcRn transgenic mice for treatment of NHL

**DOI:** 10.1101/2023.07.12.548667

**Authors:** Roman Generalov, Elisa Fiorito, Stian Foss, Veronique Pascal, Helen Heyerdahl, Ada H. V. Repetto-Llamazares, Jan Terje Andersen, Geir E. Tjønnfjord, Sigrid S. Skånland, Jostein Dahle

## Abstract

There is an unmet medical need for new therapeutic approaches and targets for patients with non-Hodgkin lymphoma (NHL) who relapse or are refractory to anti-CD20 immunotherapy. Therefore, we developed a humanized IgG_1_ antibody targeting CD37, which was tailored to be afucosylated for enhanced antibody-dependent cellular cytotoxicity (ADCC) (NNV024). In line with this, NNV024 induced three-fold more potent ADCC activity against patient-derived chronic lymphocytic leukemia (CLL) cells compared with anti-CD20 obinutuzumab. Moreover, NNV024 showed 2-fold higher ADCC activity than anti-CD20 rituximab and a recombinant version of DuoHexaBody-CD37 against both NHL and CLL cells. Survival was significantly longer after NNV024 treatment than with obinutuzumab in a mouse model. In addition, NNV024 showed a favourable plasma half-life in human FcRn transgenic mice of about 9-days, which was 2-fold longer than that of obinutuzumab and DuoHexaBody-CD37. These results warrant the further development of NNV024 as a treatment for NHL.

## INTRODUCTION

Non-Hodgkin lymphoma (NHL) is a heterogeneous group of cancers involving both B-cell and T-cell types of NHL, and indolent and aggressive NHL. The standard of care for B-cell NHL includes immunotherapy with anti-CD20 monoclonal antibodies (mAbs) alone or in combination with chemotherapy, anti-CD20 radioimmunotherapy, and small-molecule inhibitors (1, 2). Recently, new drug types, such as CAR-T cells, bispecific mAbs, and new combination treatments have been introduced, all targeting CD20 or CD19 (3–7). However, many patients ultimately relapse and become resistant to treatment, highlighting a substantial unmet need for alternative therapeutic targets and strategies (8). In recent years, CD37 has gained substantial attention as a promising therapeutic target for NHL treatment (9–11).

CD37 is a highly glycosylated transmembrane protein of the tetraspanin protein family that is expressed predominantly on mature B cells and has minimal or no expression on other hematopoietic cells (12, 13), except for mast cells (14). CD37, as part of tetraspanin-enriched microdomains (TEMs), together with other tetraspanins and integrins, is involved in paving the B-cell plasma membrane landscape (15, 16). The N-terminal domain of CD37 contains an intracellular ITIM-like motif capable of transducing pro-apoptotic signalling via the PI3K/AKT pathway, whereas the C-terminal domain carries ITAM-like motifs that mediate pro-survival signalling via the same pathway (17). A functional complex of CD37, SOCS3, and IL-6 receptor modulates cytokine signalling (16, 18). Additional functions of CD37 have been reviewed previously (17).

CD37 is highly expressed in malignant B-cells (19–21). Therapeutic anti-CD37 proteins include antibody drug conjugates (ADCs) (IMGN529, AGS67E, Naratuximab Emtansine) (22–24), a radioimmunoconjugate (RIT) (^177^Lu-lilotomab satetraxetan) (25, 26), an immuno-pharmaceutical (SMIP-016) (27), Fc-engineered mAbs (mAb 37.1 and DuoHexaBody-CD37) (28, 29), and CAR-T cells (CAR-37 H-L)(30). The mechanisms of action (MoA) of these therapeutic agents include direct cytotoxicity if conjugated with cytotoxic or radioactive payloads (ADC and RIT), classical FcγR-mediated effector functions such as ADCC (Antibody Dependent Cellular Cytotoxicity) for SMIP-016 and mAb 37.1, ADCP (Antibody Dependent Cellular Phagocytosis), CDC (Complement Dependent Cytotoxicity) for DuoHexaBody-CD37 (DHXBD37), and T-cell mediated cytotoxicity for CAR-37 H-L. The optimal MoA for the treatment of B-NHL is not clear, but ADCC and ADCP have both been utilized in successful marketed drugs, such as anti-CD20 mAbs obinutuzumab and rituximab, respectively, for the treatment of NHL (31).

Here, we report the generation of a panel of CD37-targeting humanized IgG1 mAbs, including NNV024, which is an afucosylated variant with enhanced ADCC activity. The mechanisms of action of the candidates were characterized in vitro and benchmarked against rituximab, obinutuzumab, and recombinant DHXBD37. The pharmacokinetics of NNV024 were found to be better than those of the benchmark antibodies, and in a mouse therapy study, NNV024 was superior to obinutuzumab.

## MATERIALS AND METHODS

### Antibodies

Humanized NNV mAb variants were generated by GenScript. NNV023, NNV024, NNV025, and anti CD37 Duobodies were produced by Evitria (Zurich, Switzerland). NNV024 was produced using GlymaxX® (ProBioGen, Berlin, Germany) (32). NNV003 was manufactured by Nordic Nanovector. NNV003 was conjugated to p-SCN-Bn-DOTA (NNV003-DOTA) and radiolabelled with ^177^Lu, as described previously (21). DHXBD37 was prepared as previously described (33). Rituximab and obinutuzumab were obtained from commercial sources. Recombinant versions of ofatumumab and ofatumumab P329A were produced in Expi293 cells at The Laboratory of Adaptive Immunity and Homeostasis (Oslo University Hospital, Norway).

### Cell cultures

Daudi, Ramos, Raji, Rec-1, and REH cells were obtained from ATCC (Manassas, VA, USA). U2932, SU-DHL-4, SU-DHL-6, WSU-DLCL-2, and Granta-519 cells were provided by the University Medical Center, Groningen. DOHH-2 cells were obtained from DSMZ (Braunschweig, Germany). Authentication of the cell lines was confirmed by Eurofins Genomics Europe Applied Genomics GmbH (Ebersberg, Germany). All cell lines were cultured at 37 °C in a humidified 5 % CO_2_ incubator using RPMI 1640 GlutaMAX (Gibco, Waltham, MA, USA) cell culture medium, except for Granta-519, which was cultured in DMEM high glucose (Gibco). Cell growth media were supplemented with 10 % Fetal Bovine Serum and 1 % penicillin-streptomycin (Gibco).

Peripheral blood mononuclear cells (PBMCs) were isolated from the blood of chronic lymphocytic leukemia (CLL) patients collected at Oslo University Hospital (OUS)after informed consent, and were prepared as described in (34). The patients were treatment naïve and the CLL cells presented mutated IGVH. CD37 and CD20 expression were confirmed by flow cytometry as described in the next sections.

### In silico immunogenicity prediction tools

In silico MHC class II-peptide binding prediction analysis of peptides derived from the light (LC) and heavy (HC) chain sequences of selected candidates was performed using the *NetMHCIIpan-3.1* algorithm by EIR Sciences APS (35). De-immunization of the LC sequence spanning the mV_L_-hC_L_ region was performed in silico by introducing all-natural amino acids at the positions of interest. Conservation scores were calculated using the PSI-BLAST algorithm, as described previously (36).

### Competition binding assay

Binding of the mAb variants to Ramos cells was assessed by a competition binding assay, as described previously (37). Ramos cells were washed twice and 8.0 x 10^5^ were aliquoted into thirteen 12×75 mm glass reagent tubes (VWR) for each test antibody. The cells were maintained on ice during the experiment. The test antibodies were dispensed to the cells at concentrations ranging 0.001 - 1000 nM. NNV003-DOTA was radiolabelled with ^177^LuCl_3_ (ITG Isotope Technologies Garching GmbH, Germany) as described in (21, 38) and added to each tube to a final concentration of 2.7 nM. The tubes were placed on the Wizard2 3470 Gamma Counter and assayed for 1 min. The duration of incubation with the radiolabelled ligand was 4 hrs at 4°C, 350 RPM on orbital shaker. The cells were washed three times with 0.5 mL of ice-cold PBS with 0.5% BSA. After the last wash, the tubes containing the cell pellets were assayed on the gamma counter using the same counting protocol. The IC_50_ and the affinity parameter 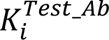 was calculated for each corresponding test antibody using One Site Competition option in the build-in “Simple Ligand Binding” macros of SigmaPlot 12.0 (39).

### ADCC and ADCP reporter assays

The assays were conducted following the protocols from Promega Biotech AB (Nacka, Sweden) and described previously (40). The assays were conducted following the protocols recommended in the relevant Promega kits from Promega Biotech AB (Nacka, Sweden). In short, target cells (T) were harvested, washed three times in PBS with 0.5% BSA, resuspended in RPMI supplemented with 4% low IgG serum (assay buffer) to concentration 1.0 × 10^6^ cells/mL and seeded (2.5×10^4^ cells/well) in a white flat bottom 96 well plate (165306, Thermo Fisher). Right after plating the cells, the plates were conditioned to ambient temperature, 21°C (RT). Serial dilutions (0.001 – 1 µg/mL in assay buffer) of the test antibodies NNV023, NNV024, obinutuzumab, rituximab, and DuoHexabody-CD37 were added to the wells and incubated for 30 min at 21°C to induce opsonization. Some wells with target cells were left without addition of any Ab. The Promega effector Jurkat cells (E) recombinantly modified to stably express either FcγRIIIa-V158 (ADCC), FcγRIIa-H131 or FcγRIIa-R131 (both ADCP) receptors were thawed, resuspended in the assay buffer and added to the target cells in proportion 3:1 (E:T). The system was incubated for 16-18 hours followed by addition (equivolumetric ratio) of Bio-Glo Luciferase Assay Reagent (Promega) at RT. The luminescence read out was performed using a Spark microplate reader (TECAN, Switzerland). Normalized induction of ADCC or ADCP was calculated by dividing the luminescence values of the treated cells by the mean luminescence of untreated (Ab-free) cells.

### Complement-dependent cytotoxicity (CDC) assay

The target cells were prepared as outlined in ADCC and ADCP assays. Serially diluted (0.01-10 µg/mL) test antibodies NNV023, NNV024, NNV025, obinutuzumab, rituximab, DuoHexabody-CD37, ofatumumab WT (positive control) or ofatumumab P329A (negative control) were added to the cells at RT. A well of untreated cells (Ab-free control) and no-cell wells (blank) were included in each plate. The plates were incubated for 30 minutes in a humidified incubator (37 °C, 5% CO_2_) to promote cell opsonization. Human serum complement (CTS-006, Creative Biolabs) was added (12.5% final concentration) to all wells and the plates were placed in the incubator for two hours. Then, AlamarBlue (Invitrogen/Life Technologies) was added to each well 1:10 v/v. After 16-24 hours in 37°C, 5% CO_2_ incubator, the fluorescence of each well was recorded (λ_exc_: 544 nm, λ_em_: 590 nm) using Fluoroskan Ascent FL plate reader (Thermo Electron Corporation). The mean of triplicate measurements of the cell-containing wells were divided by the mean of the blank wells to normalize signal to baseline. Then, the normalized mean fluorescence of each treated well was divided by the normalized mean fluorescence of untreated wells (Ab-free control). The results were expressed as percentage of the untreated control.

### Binding to FcRn and FcγRs

Ninety-six-well EIA/RIA assay microplates (#3590; Corning) were coated with 100 µL/well of serially diluted test mAbs (0.45 – 10,000 ng/mL) in PBS overnight at 4°C. After blocking and washing the plates as described previously (41), the pre-formed complexes of 250 ng/mL biotinylated FcγRIIa-H131, FcγRIIa-R131, FcγRIIb, FcγRIIIa-V158, FcγRIIIa-F158, or FcγRIIIb (all from Sino Biological, Germany) and alkaline phosphatase (AP)-conjugated streptavidin (Roche Diagnostics, UK) were added to the wells and incubated for 1 h at RT. High-affinity FcγRI (Sino Biological) and AP-streptavidin were added separately. Bound receptors were visualized by the addition of 1 mg/mL (final concentration) p-nitrophenylphosphate in diethanolamine buffer. The plates were incubated for 15-20 minutes before absorption at 405 nm was measured using a TECAN Sunrise plate reader (TECAN, Switzerland).

Binding of mAbs to human FcRn was performed by measuring surface plasmon resonance (SPR) using a Biacore T200 instrument (GE Healthcare, UK), as described previously (42).

### In vivo PK studies

Male hemizygous human FcRn transgenic mice (B6.Cg-Fcgrt^tm1Dcr^ Tg(FCGRT)32Dcr/DcrJ) (Tg32 hemizygous) that are knockout for the mouse FcRn heavy chain and express the genomic transgene of the human FcRn heavy chain under the control of the human FcRn promoter were used to study the plasma half-life of mAbs (Jackson laboratories (JAX), USA). The in vivo part of the study was performed by JAX Services in an AAALAC^1^-accredited animal facility under IACUC^2^ review. Mice were kept under pathogen-free conditions in a 12-hour light/dark cycle, with access *ad libitum* to food and water. Temperature, humidity, and airflow were monitored continuously.

In the first experiment, mice were administered 5 mg/kg NNV025, NNV023, NNV024, obinutuzumab, or DHXBD37 on day 0. Blood samples were collected into tubes containing 1 % tripotassium-ethylene diamine-tetraacetic-acid (K3-EDTA) and centrifuged to isolate plasma. The plasma was diluted 1:10 in glycerol/PBS solution and stored at -20°C until analysis at Oslo University Hospital. Quantification of mAbs was performed using an anti-human Fc sandwich ELISA as described previously (43).

The absorbance values obtained from the ELISA analysis were interpolated to standard curves covering the antibody ranges using Graphpad Prism v.8.3.0. The plasma concentration of each antibody samples at each time point was then calculated by multiplying with the sample dilution factor (Microsoft Excel). For the evaluation of plasma clearance, the first concentration measured (day 1) was normalized to 100% and remaining data points were plotted as percent antibody remaining in plasma. Data points from the β-elimination phase were then used to calculate the plasma half-life using the formula:

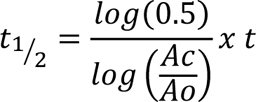

where *t_1/2_* is the half-life, *A_c_* is the amount of antibody remaining, and *A_0_* is the amount of antibody at day 1 and *t* is the elapsed time (44).

The antibody concentrations in plasma determined by ELISA were then fitted to a non-compartmental (NCA) PK model using the gPKPDsim PK add-on for MatLAb (45). Briefly, the antibody concentrations were plotted in Microsoft Excel and loaded into gPKPDsim before both half-life and PK parameters were calculated using predefined formulas (45).

### In vivo proof-of-concept studies

Efficacy studies were blinded and conducted at ArcticLAS (Reykjavik, Iceland) under animal license number 2019-03-04. ArcticLAS is authorized by the Food and Veterinary Authority of Iceland (MAST).

Mice were housed under pathogen-free conditions in a 12-hour light/dark cycle with access *ad libitum* to food and water. Temperature, humidity, and airflow were monitored continuously.

In the first study, 70 female CB17-SCID mice, 6-7 weeks of age (Envigo, France), were weighed, earmarked, and randomized into seven study groups, with 10 animals per group. All mice received 1×10^7^ Daudi cells intravenously on day -1. The next day, the mice were treated with 100 µg NNV024 or obinutuzumab. The mAbs were administered to the animals as either one injection (on day 0), three injections (days 0, 4, and 9), or six injections a course (days 0, 4, 9, 12, 16, and 19). The Animals in the control group were treated with 0.9 % NaCl (days 0, 4, and 9).

In the second study, 50 female CB17-SCID mice (Envigo) were prepared as described above and treated with a single dose of 10 or 50 µg NNV024, 10, or 50 µg obinutuzumab, or 0.9 % NaCl.

In both experiments, animals were euthanized if one or more humane endpoints were reached: hind leg paralysis, overall weight loss of ≥20 %, or signs of substantial discomfort. At termination, all animals were necropsied, the organs were investigated for signs of tumors, and selected organs were harvested for histopathological evaluation of microscopic tumor infiltration.

### Statistical methods

Kinetic rate constants, binding affinity and plasma half-lives were compared using Student t-test.

Differences in *in vivo* antibody clearance and plasma half-lives were also analyzed in GraphPad Prism v.8.3.0 using repeated measurements two-way ANOVA with Geisser-Greenhouse correction.

Survival analysis was performed using the log-rank test (Mantel-Cox Test), and pairwise comparison was performed using the Holm-Sidak method (α= 0.05). GraphPad Prism v.8.3.0 was used for calculations of median survival and p-values for each treatment group.

## RESULTS

### Selection and development of humanized Ab lead candidate

A panel of humanized fragment antigen-binding (Fab) variants of the anti-CD37 lilotomab mAb was generated by the direct grafting of complementary determining regions to the human IgG_1_ acceptor sequences. A phage library generated to exploit 14 putative back mutations was used for affinity maturation via two rounds of panning against CD37-expressing Ramos cells. The Fab sequences carrying the lowest number of back mutations were selected for further development and converted into full-length antibodies using a human IgG_1_ backbone. The hIgG_1_ mAbs all exhibited a higher binding affinity to CD37 than chimeric NNV003, but there was no major difference between the variants (Table 1).

**Table 1.** Immunogenicity risk score and binding affinity to CD37 of the selected humanized mAb variants

| Designation | Immunogenicity Risk Score<sup>1</sup> |  |  | Binding Affinity<sup>5</sup><br>K<sub>D</sub> ± SE, (nM) |
| --- | --- | --- | --- | --- |
|  | HC | LC | Combined |  |
| Lilotomab | 20.5 | 15.8 | 36.4 | 1.6 ± 1<sup>6</sup> |
| NNV003 | 7.3 | 15.8 | 23.1 | 1.0 ± 0.07<sup>7</sup> |
| NNV037<sup>2</sup> | 8.0 | 9.2 | 17.2 | 0.53 ± 0.09 |
| NNV028<sup>2</sup> | 7.5 | 9.2 | 16.7 | 0.33 ± 0.06 |
| NNV036<sup>2</sup> | 5.3 | 9.8 | 15.2 | 0.44 ± 0.04 |
| NNV035<sup>2</sup> | 7.2 | 7.9 | 15.1 | 0.42 ± 0.06 |
| NNV026<sup>2</sup> | 4.5 | 9.2 | 13.6 | 0.54 ± 0.07 |
| NNV027<sup>2</sup> | 4.8 | 8.5 | 13.3 | 0.53 ± 0.10 |
| NNV025<sup>3</sup> | 4.5 | 7.9 | 12.4 | 0.53 ± 0.06 |
| NNV023<sup>4</sup> | 4.5 | 5.6 | 10.0 | 0.57 ± 0.08 |
| Rituximab | 4.7 | 12.6 | 17.4 | Not tested |

<sup>1</sup> World average population. <sup>2</sup>First set of selected candidates. <sup>3</sup>NNV025: NNV026 HC + NNV035 LC sequences. <sup>4</sup>NNV023: NNV026 HC + NNV035-V110D LC sequences. HC: Heavy chain. LC: Light chain. <sup>5</sup>Cell-based competitive binding assay using CD37+ Ramos cells. <sup>6</sup>From Table 5 in (25). <sup>7</sup>Average value from Table 1S in (38).

Immunogenicity risk scores (IRS) showed that replacing the murine constant regions with humans decreased IRS by reducing the number of T cell neo-epitopes and/or their promiscuity (Table 1 and Supplementary Figure 1). The NNV035 light chain (LC) and NNV026 heavy chain (HC) sequences were selected for the generation of NNV025, the variant with the lowest combined IRS. In silico MHC class II binding peptide mapping identified a promiscuous neo-epitope, which was not observed in lilotomab, spanning the mV_L_-hC_L_ region of the NNV003 and NNV025 sequences (Supplementary Figure 1). This epitope has also been previously identified in rituximab (46). In silico de-immunization of the NNV035_LC sequence was performed to reduce the promiscuity of the predicted epitope spanning positions 105–115 by introducing the V110D amino acid substitution without impacting CD37 binding (Supplementary Figure 1). The NNV026 HC and NNV035_V110D LC sequences were selected to generate NNV023, which was predicted to have the lowest IRS. Furthermore, an afucosylated version of NNV023, NNV024, was manufactured by recombinant co-expression of NNV023 sequences with the bacterial oxidoreductase GDP-6-deoxy-D-lyxo-4-hexulose reductase to enhance its ADCC effector function.

### In vitro binding to FcγRs

ADCC activity is dependent on binding to FcγRIIIa^V158/F158^ expressed on NK cells, macrophages, and monocytes (47) whereas ADCP activity is dependent on binding to FcγRIIa^H131/R131^, FcγRIIIb, FcγRI, and FcγRIIb. Therefore, the binding affinities of NNV025, NNV023, NNV024, obinutuzumab, and DHXBD37 to these receptors were measured. As expected, glycan -engineering increased the binding affinity of NNV024 to FcγRIIIa and FcγRIIIb (Figure 1 A, B, and F). As compared to NNV023, the affinity of NNV024 to F158 and V158 allelic variants of FcγRIIIa were 4.4-fold (from 460.8 to 104.1 ng/mL) and 2.7-fold higher (from 860.7 to 332.4 ng/mL), respectively (Supplementary Table 1). The affinity of NNV024 to FcγRIIIa was almost three times stronger than that of obinutuzumab for both F158 and V158 allelic variants in the ELISA format. Afucosylation also increased affinity for FcγRIIIb by 2.8-fold. Therefore, enhanced induction of ADCC should be expected for NNV024 compared to the other tested mAbs. The afucosylation did not affect the binding affinity of NNV024 to the ADCP associated receptors, FcγRIIa^H131/R131^, FcγRIIb, and FcγRI (Figure 1 C, D, E, and G; Supplementary Table 1). Obinutuzumab and DHXBD37 exhibited low binding affinities to FcγRIIa, FcγRIIb, and FcγRI in ELISA settings.

**Figure 1.**
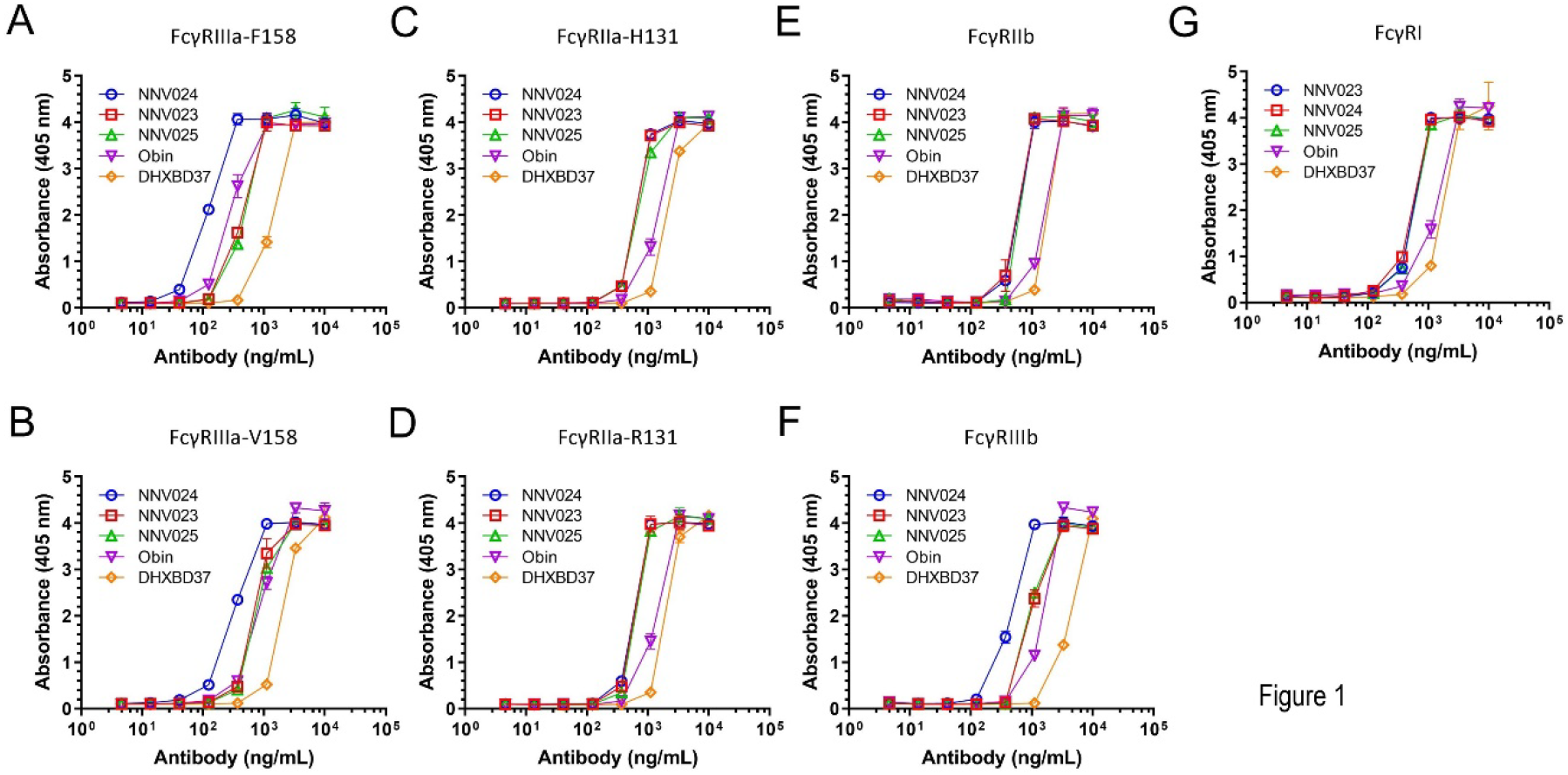
Binding to human FcγRs. The results of ELISA showing concentration-dependent binding of the Abs (4.5 – 10.000 ng/mL) to biotinylated human FcγRIIIa^V158^, FcγRIIIa^F158^, FcγRIIa^H131^, FcγRIIa^R131^, FcγRIIb, FcγRIIIb, and FcγRI. Data shown as mean ± SD. Obin – obinutuzumab; DHXBD37 – DuoHexaBody-CD37. The experiment was performed once with two replicates.

### Induction of ADCC, ADCP and CDC in NHL cell lines

In vitro ADCC potency was investigated in a panel of 11 B-NHL cell lines of different subtypes and in patient-derived CLL cells, using a reporter assay with cells expressing recombinant FcγRIIIa-158F. NNV024 induced stronger ADCC activation than NNV023, rituximab, and DHXBD37 (Figure 2 A), as expected from the in vitro binding data (Figure 1). However, both NNV024 and obinutuzumab fully activated ADCC. The degree of ADCC induction depends on the target cells. EC_50_ represents Half Maximal Effective Concentration or the potency of a drug. The lower the EC_50_ value, the higher potency is, the less of the drug is needed to reach the therapeutic dose. EC_50_ of NNV024 prevailed over obinutuzumab in all cell lines except for Raji, U2932 and Rec-1, where they performed equally.

**Figure 2.**
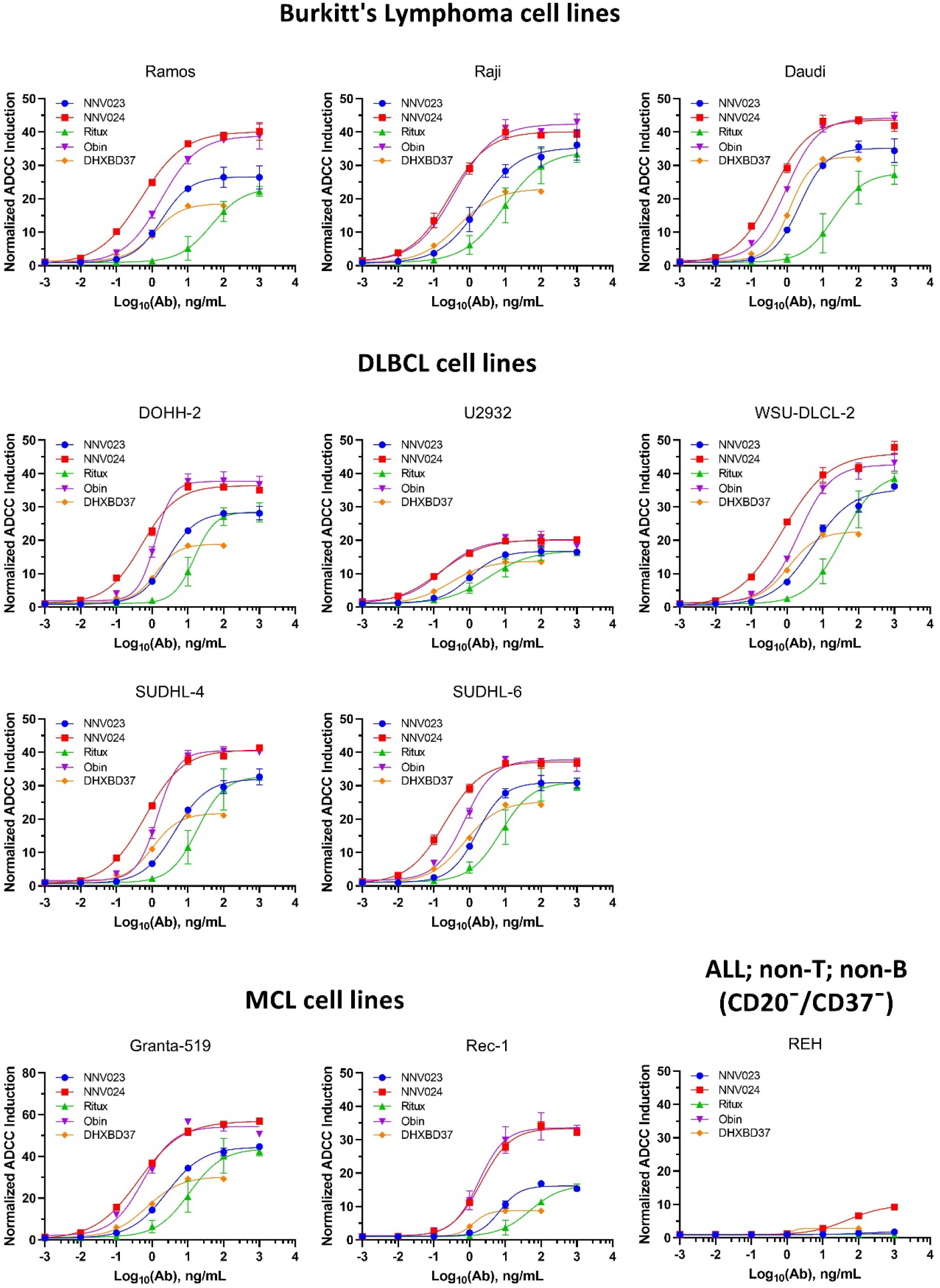
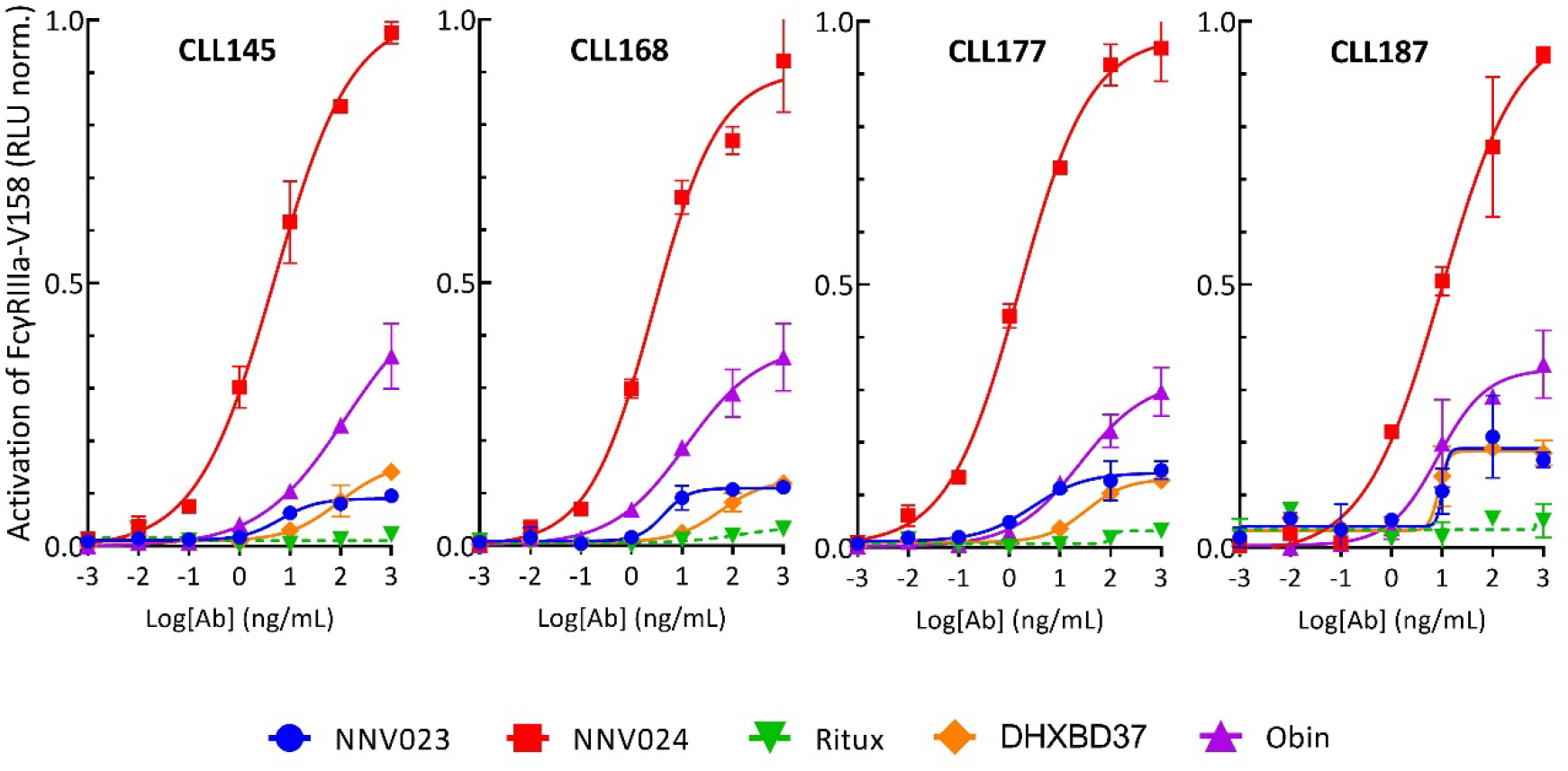
ADCC induction. Normalized ADCC induction assessed with ADCC FcγRIIIa-158F reporter assay in cell lines (A) and treatment naïve CLL patient samples (B). Abbreviations: DLBCL – Diffused Large B-Cell Lymphoma; MCL – Mantle Cell Lymphoma; CD20^-^/CD37^-^ CTRL – a control cell line (REH) that express neither CD20 nor CD37; Ritux – rituximab; Obin – obinutuzumab; DHXBD37 – DuoHexaBody-CD37. The figure shows one representative experiment of two with three replicates.

ADCC induction of NNV023 was, on average, 40 % lower than that of NNV024 and 40 % above rituximab (Supplementary Figure 3).

NNV023 and NNV024 did not induce ADCP or CDC in any tested cell line (Supplementary Figure 4 and 5). In contrast, the ADCP and CDC effects of DHXBD37, and rituximab were pronounced. Obinutuzumab had a low CDC response, but high ADCP. For CDC, ofatumumab was also included as a positive control. CDC data were in line with previously reported observations for DHXBD37 (29), ofatumumab, and rituximab (29, 48). It was found no correlation between the expression levels of CD37 or CD20 and *in vitro* ADCC parameters such as Emax, EC_50_, AUC (Supplementary Figures 6 and 7).

### Induction of ADCC in patient derived CLL cells

NNV024 was the strongest inducer of ADCC in patient derived CLL cells (Figure 2B). The ADCC signaling induced by 1 µg/mL NNV024 in all four CLL patient samples was approximately 3-fold higher than that induced by 1 µg/mL obinutuzumab. Fully fucosylated NNV023 induced ADCC on par with DHXBD37, while rituximab induced the lowest ADCC in CLL patient samples. The CD37 expression was strong in the patient derived CLL cells (Supplementary Figure 8).

### In vitro binding to hFcRn

Binding to the human neonatal Fc receptor (hFcRn) was tested because this receptor is involved in the extension of the half-life of IgG by rescuing IgG from proteolytic degradation in lysosomes. Binding to hFcRn was tested by SPR under both acidic (pH 6.0) and neutral (pH 7.4) conditions to mimic the endosomal and extracellular milieu, respectively. All mAbs bound to hFcRn under acidic conditions, but at neutral pH, the interaction was very weak (Supplementary Figure 2). The kinetic rate constants and binding affinity for hFcRn by SPR under acidic conditions for the NNV mAbs were not significantly different from those of obinutuzumab and DHXBD37 (Table 2).

**Table 2.** Kinetic rate constants and binding affinity of mAbs to hFcRn at pH 6.0 measured by SPR.

| mAb | K<sub>a</sub> (1/Ms)<sup>1</sup> | K<sub>d</sub> (1/s) | K<sub>D</sub> (M) |
| --- | --- | --- | --- |
| NNV025 | 4.849x10<sup>5</sup> | 0.1020 | 2.103x10<sup>-7</sup> |
| NNV023 | 4.170x10<sup>5</sup> | 0.1060 | 2.542x10<sup>-7</sup> |
| NNV024 | 4.591x10<sup>5</sup> | 0.1039 | 2.262x10<sup>-7</sup> |
| Obinutuzumab | 4.776x10<sup>5</sup> | 0.0917 | 1.920x10<sup>-7</sup> |
| DuoHexabody-CD37 | 4.641x10<sup>5</sup> | 0.1338 | 2.883x10<sup>-7</sup> |

<sup>1</sup>Kinetic rate constants were determined using a first-order (1:1) Langmuir biomolecular interaction model.

### Pharmacokinetic characterization of mAb variants in mice expressing human FcRn

If binding to hFcRn is important for the pharmacokinetic properties of the test mAbs, one would expect a similar plasma half-life for the mAbs in transgenic mice expressing human FcRn. However, the plasma half-life of NNV023 was significantly longer than that of NNV025 in the absence of competing IgG, indicating that the amino acid substitution (V110D) introduced to reduce the risk of immunogenicity also increased the plasma half-life by 4.6 days (p < 0.001, Figure 3 and Supplementary Table 2). In contrast, afucosylation of NNV023 led to a more than 3-days reduction in the plasma half life of NNV024. The plasma half-lives of obinutuzumab (4.4 ± 1.3 days) and DHXBD37 (4.1 ± 1.2 days) were statistically significantly shorter than for NNV023 (12 ± 2.4 days) and NNV024 (8.9 ± 0.8 days, p< 0.01, t-test, Supplementary Table 2).

**Figure 3.**
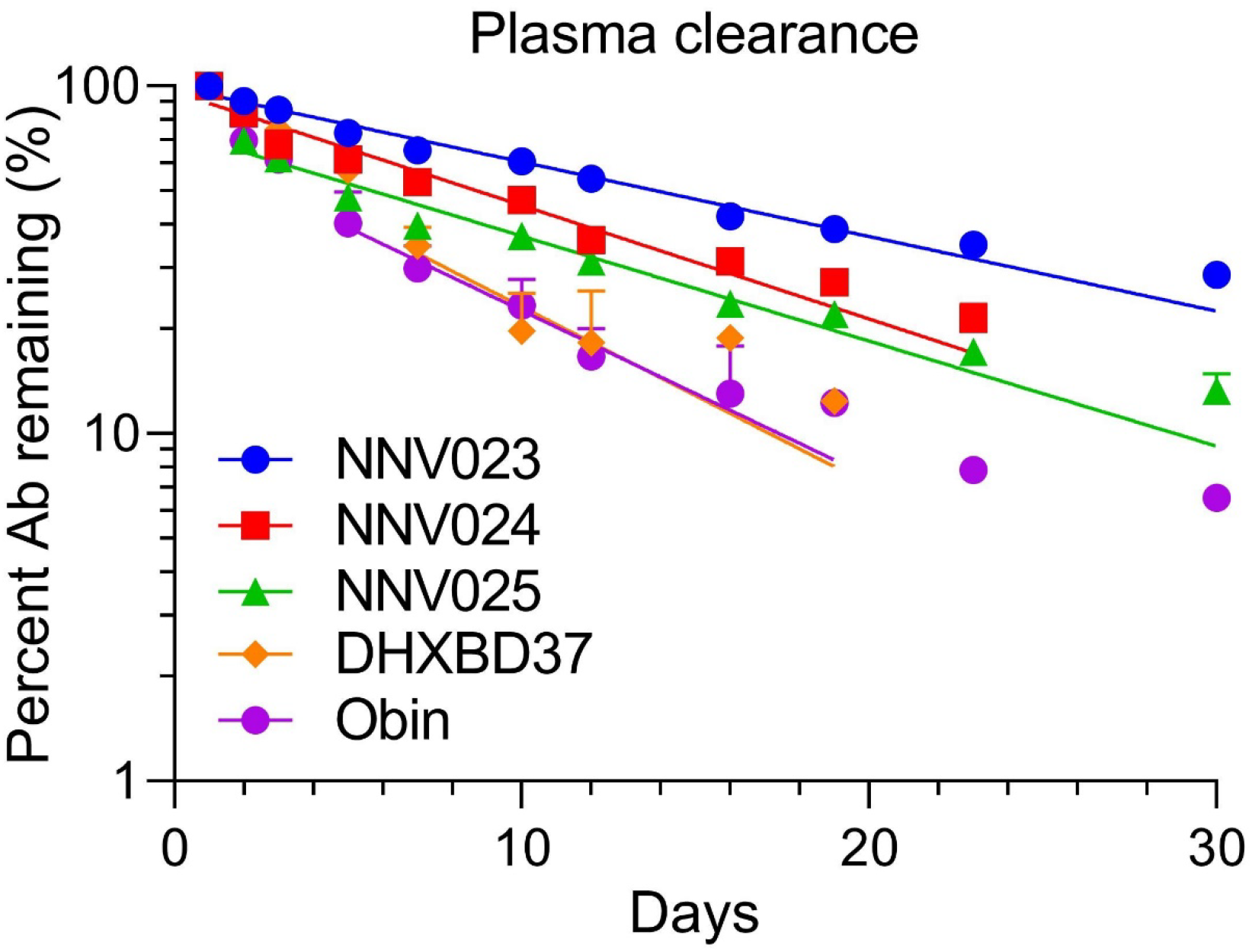
Plasma clearance. Plasma clearance of NNV023 (n=5), NNV025 (n=4), NNV024 (n=4), obinutuzumab (n=5) and DuoHexabody-CD37 (n=5) in Tg32 hemizygous mice. Error bars = SD. Abbreviations: Obin – obinutuzumab, DHXBD37 – DuoHexaBody-CD37.

### In vivo proof-of-concept studies

NNV024 was equally effective in treating disseminated B-cell malignancy as obinutuzumab at the tested dose of 100 µg, and an increased number of injections did not improve survival (Figure 4 A). Obinutuzumab was used as the comparator Ab because it is an afucosylated Ab with the same mechanism of action as NNV024 and it is approved for treatment of B-NHL. This was considered more relevant that using DHXBD37 which has enhanced CDC activity and is in clinical development. The median survival of the mAb treatment groups was in the range of 77-90 days, whereas that of the control group was 24 days (Supplementary Table 3). All treatment groups had significantly longer survival than the control group (p<0.0001), and there was no statistically significant difference in survival between the treatment groups (p>0.05, Mantel-Cox Test) (Supplementary Table 3).

**Figure 4.**
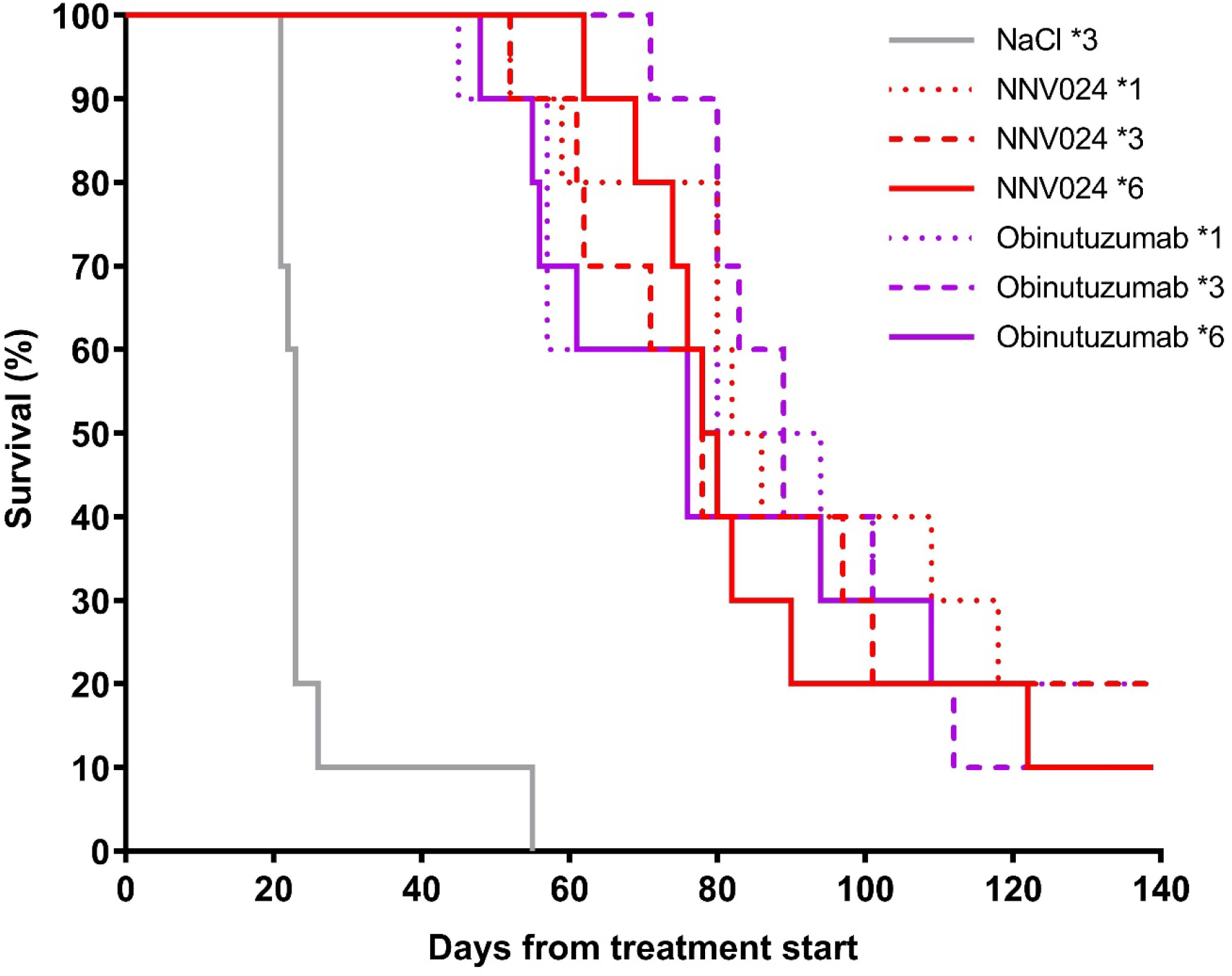
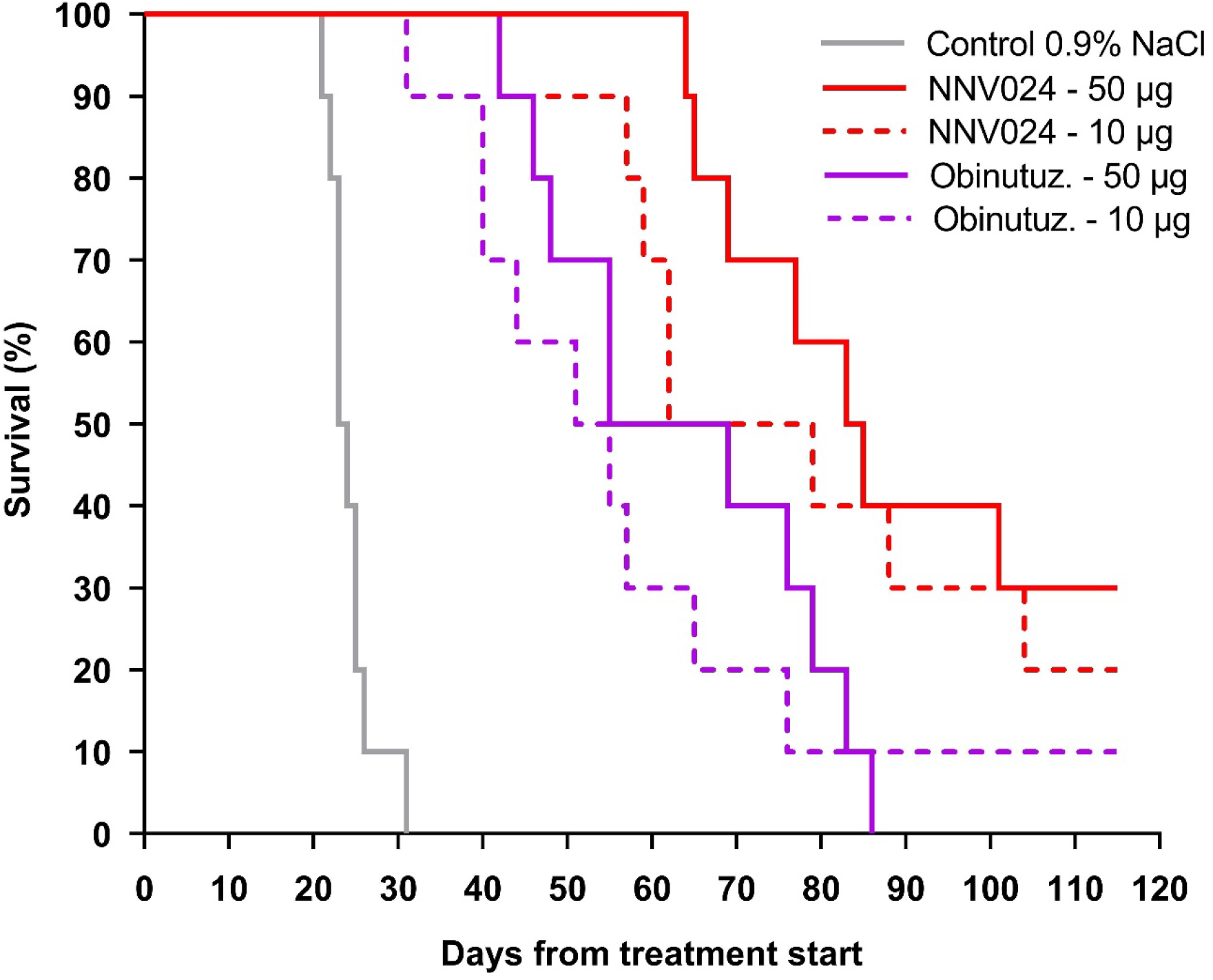
Proof of concept studies. A) Survival of CB17-SCID mice with i.v. injected Daudi cells treated with 1, 3 or 6 injections of 100 μg NNV024 or obinutuzumab or 100 µl 0.9 % NaCl. All treatment groups had statistically significant better survival rates than the control group (p < 0.0001). There was no statistically significant difference in survival between treatment groups (p>0.05, Mantel-Cox Test). B) Survival of CB17-SCID mice with i.v. injected Daudi cells (1×10^7^ cells) and treated with 10 or 50 μg of NNV024 or obinutuzumab or 100 µl 0.9 % NaCl. All treatment groups (NNV024 or obinutuzumab) had statistically significant better survival than the control group, p < 0.0001 (Log-rank/Mantel-Cox Test). Group NNV024 50 µg had a statistically significant better survival rate (Holm-Sidak method, α=0.05, p < 0.05) than both of the obinutuzumab groups (10 µg and 50 µg). Differences in survival of the NNV024 10 µg group compared to the NNV024 50 µg and both obinutuzumab groups were not statistically significant (p > 0.05).

The control group started losing weight 18 days after the start of treatment (Supplementary Figure 9), and all mice, except one, were euthanized 8 days later with hind leg paralysis. The average body weights of the mice treated once with obinutuzumab started to decrease 42 days after start of treatment while it started to decrease at 76 days for mice treated once with NNV024. All mice in the control group had visible tumors in the kidneys and ovaries, whereas only four mice treated with obinutuzumab and no mice treated with NNV024 had visible tumors in the kidneys (Supplementary Figures 11 and 12 shows representative images of Daudi-induced ovarian and kidney).

It was suspected that the 100 μg dose saturated the therapeutic capacity of the tested mAbs; therefore, a second efficacy study with lower doses was initiated. In this study, mice treated once with 50 μg NNV024 had a significantly longer survival than mice treated with 10 or 50 μg obinutuzumab (p = 0.012 and p = 0.016, respectively) (Figure 4 B, Supplementary Table 4). The median survival of mice treated with 50 μg NNV024 was 84 days, whereas mice treated with 10 or 50 μg obinutuzumab had a median survival of 53 and 62 days, respectively. All treatments with NNV024 and obinutuzumab resulted in a significantly prolonged survival compared to the control group, which had a median survival of 23.5 days (p < 0.0001). Mice treated with 10 μg NNV024 had a median survival of 70.5 days, which was longer than that of mice treated with obinutuzumab; however, the difference was not statistically significant.

The control group started losing weight after 15 days (Supplementary Figure 10), and all mice were euthanized 15 days later due to hind leg paralysis. The average body weight started to decrease on days 30 and 34 for the mice treated with 10 and 50 μg of obinutuzumab, respectively, while it started to decrease at days 38 and 56 for the mice treated with 10 and 50 μg of NNV024, respectively. All mice in the control group had visible tumors in the kidneys and ovaries at necropsy, whereas only two and nine out of 40 mice in all four mAb treatment groups had visible tumors in the kidneys and ovaries, respectively; however, there were no significant differences between the treated groups.

## DISCUSSION

The treatment landscape for B-cell malignancies has evolved rapidly in recent years with the introduction of new drugs such as CAR-T cells, bispecific Abs, and novel combination treatments (3–7). However, traditional targeted therapies for NHL and many new treatments are mainly based on the CD20 antigen. Therefore, resistance to CD20-based therapies remains an unmet medical need for many patients with NHL. Furthermore, some new treatment options are associated with a substantial risk of cytokine release syndrome and other toxicities (3–7). Consequently, treatments against new targets are required. CD37 is an alternative B-NHL target that is highly expressed on mature B cells and most B-cell lymphomas (21) and has recently been clinically validated (24, 26, 28). With this in mind, we developed NNV024, a monoclonal humanized anti-CD37 mAb with optimized predicted immunogenicity and enhanced affinity to FcγRs, as an alternative and potentially safer treatment option for CD37-positive B-NHLs. NNV024 showed a significantly improved therapeutic effect compared to obinutuzumab in a B-NHL mouse model. The in vivo data were supported by both stronger binding to FcγRIIIa and FcγRIIIb and more potent activation of ADCC by NNV024 than by obinutuzumab. These encouraging preclinical results show that NNV024 is a highly promising mAb for the treatment of B-cell malignancies.

Median survival was prolonged up to 40 % with a single treatment of NNV024 compared to obinutuzumab. Increasing the dose from 50 to 100 μg did not improve NNV024 efficacy; median survival was 84 and 85 days, respectively, but it improved the efficacy of obinutuzumab, as the median survival increased from 62 to 88 days. However, multiple injections did not improve the potency of either of the antibodies. This result can be partly explained by the higher ADCC effect exhibited by NNV024 in Daudi cells already at doses below 0.01 µg/mL. SCID mice have functional NK cells to which human antibodies can bind with high affinity and thus induce ADCC in a mouse model (49).

Another parameter that might explain the observed difference in in vivo efficacy is residence time. In Tg32 transgenic mice expressing human FcRn, NNV024 has an almost two-fold greater half-life in plasma than obinutuzumab, 8.9 ± 0.8 days versus 4.4 ± 1.3 days, respectively. Therefore, the extended plasma half-life of NNV024 may increase NNV024 exposure at the target sites. While the SCID mice used in the in vivo efficacy study did not express human FcRn, it can be speculated that differences in plasma half-life could account for some of the observed differences in efficacy. It is worth noting that the difference in CD37 expression level compared to CD20 was probably not the underlying mechanism promoting NNV024 impact, as the expression of the two antigens on Daudi cells has been shown to be similar (21). Therefore, we propose that it is the combination of a higher ADCC efficacy with a prolonged residence time that explains the increased potency exhibited by NNV024 compared to obinutuzumab in vivo. Cross-reactivity studies conducted for NNV024 (unpublished data) and obinutuzumab (50) excluded any potential cross-reactivity with either CD20 or CD37 murine analogs from the reported results.

The plasma half-life of NNV023 was 62 % longer than that of NNV025 in a Tg32 hemizygous mouse model. We attribute this to the V110D mutation in the light chain of NNV023, which was the only difference between the two antibodies. No difference was found in binding to human FcRn; therefore, the enhancement in half-life cannot be explained by Abs recycling via the FcRn salvage mechanism.

Afucosylation of NNV023 to NNV024 led to a 26 % reduction in plasma half-life, from 12.0 to 8.9 days. It is widely accepted that N297-linked oligosaccharides contribute to interactions with FcγRs and C1q but not with FcRn (51). However, the complete absence of mannose or, in contrast, the presence of high mannose on the N-glycan has been shown to decrease the plasma half-life (52, 53). Furthermore, hypersialylation has been reported to increase the circulatory half-life of monoclonal antibodies (54, 55). However, in the current case, it seems that afucosylation partly counteracts the effect of the V110D mutation on plasma half-life.

Our data suggest that NNV024 could be an attractive therapeutic alternative to obinutuzumab for the treatment of NHL patients, with the possibility of reaching a similar or improved response to treatment with a lower dose of mAb. Nevertheless, although the selection of the drug candidate NNV024 integrated in silico prediction tools to reduce the predicted immunogenicity potential of lilotomab, additional biological factors such as antigen processing and T cell receptor repertoire for MHC class II/peptide complexes, as well as drug-related parameters such as post-translational modifications (e.g., glycosylation profile), structure (e.g., aggregates), and impurities (e.g., host cell protein content) can affect the clinical immunogenicity of a therapeutic protein. Development of an anti-drug antibody response could then impact the efficacy of the drug by impairing pharmacokinetics, but also impact patient safety. Conclusions regarding NNV024 immunogenicity and any impact on drug efficacy will therefore have to be addressed further in the clinic.

A recent study showed high expression of CD37 in mast cells (14) which are involved in autoimmune diseases, such as rheumatoid arthritis, multiple sclerosis, insulin-dependent diabetes mellitus, and chronic idiopathic urticaria, through different mechanisms, including the release of cytokines/chemokines to recruit and activate T cells and macrophages (56). B cells also play a central role in the pathogenesis of autoimmune diseases, as the loss of B cell tolerance results in increased serum levels of autoantibodies, enhanced effector T cell response, and tissue damage (57). Although B cell depletion can eliminate most circulating B cells in the periphery, the clinical outcomes of B cell depletion therapy for autoimmune diseases vary among individuals due to differential activation or survival signals for B cells provided by the tissue microenvironment and/or mast cells. Therefore, the depletion of both B cells and mast cells by NNV024 in autoimmune diseases could be an interesting next step in the development of NNV024.

The encouraging preclinical results shown for NNV024 warrant further exploration of this therapeutic mAb for the treatment of B-cell malignancies as well as B-cell-driven autoimmune disorders.

## ACKNOWLEDGEMENTS

The authors acknowledge the assistance of Bergthora Eiriksdottir at ArcticLAS for the in vivo proof-of-concept studies.

## Author contributions

Conceptualization, J. D., R. G., and V.P.; methodology, J.D., R.G., V.P., H.H., E.F., A.R.L., and J.T.A; investigation, R.G., E.F., S.F., V.P., H.H., and A.R.L.; patient samples, G. E. T. and S.S.S.; writing, draft preparation, review, and editing, all authors. All the authors have read and agreed to the submitted version of the manuscript.

## Data access

All relevant data are within the paper and its Supporting Information files.

## Institutional Review Board Statement

PK studies were approved by the Institutional IACUC committee. Therapy studies were approved by the Regional Committee for Medical and Health Research Ethics of South-East Norway (2016/947). The research on human blood was conducted in accordance with the Declaration of Helsinki.

^1^ The American Association for Accreditation of Laboratory Animal Care

^2^ Institutional Animal Care and Use Committee

## REFERENCES

1. Salles G, Barrett M, Foa R, Maurer J, O’Brien S, Valente N, et al. Rituximab in B-Cell Hematologic Malignancies: A Review of 20 Years of Clinical Experience. Adv Ther. 2017;34(10):2232–73.

2. Crisci S, Di Francia R, Mele S, Vitale P, Ronga G, De Filippi R, et al. Overview of Targeted Drugs for Mature B-Cell Non-hodgkin Lymphomas. Front Oncol. 2019;9:443.

3. Budde LE, Assouline S, Sehn LH, Schuster SJ, Yoon SS, Yoon DH, et al. Single-Agent Mosunetuzumab Shows Durable Complete Responses in Patients With Relapsed or Refractory B-Cell Lymphomas: Phase I Dose-Escalation Study. J Clin Oncol. 2022;40(5):481–91.

4. Hutchings M, Morschhauser F, Iacoboni G, Carlo-Stella C, Offner FC, Sureda A, et al. Glofitamab, a Novel, Bivalent CD20-Targeting T-Cell-Engaging Bispecific Antibody, Induces Durable Complete Remissions in Relapsed or Refractory B-Cell Lymphoma: A Phase I Trial. J Clin Oncol. 2021;39(18):1959–70.

5. Schuster SJ, Bishop MR, Tam CS, Waller EK, Borchmann P, McGuirk JP, et al. Tisagenlecleucel in Adult Relapsed or Refractory Diffuse Large B-Cell Lymphoma. N Engl J Med. 2019;380(1):45–56.

6. Leonard JP, Trneny M, Izutsu K, Fowler NH, Hong X, Zhu J, et al. AUGMENT: A Phase III Study of Lenalidomide Plus Rituximab Versus Placebo Plus Rituximab in Relapsed or Refractory Indolent Lymphoma. J Clin Oncol. 2019;37(14):1188–99.

7. Sehn LH, Chua N, Mayer J, Dueck G, Trneny M, Bouabdallah K, et al. Obinutuzumab plus bendamustine versus bendamustine monotherapy in patients with rituximab-refractory indolent non-Hodgkin lymphoma (GADOLIN): a randomised, controlled, open-label, multicentre, phase 3 trial. Lancet Oncol. 2016;17(8):1081–93.

8. Klener P, Klanova M. Drug Resistance in Non-Hodgkin Lymphomas. Int J Mol Sci. 2020;21(6).

9. Beckwith KA, Byrd JC, Muthusamy N. Tetraspanins as therapeutic targets in hematological malignancy: a concise review. Front Physiol. 2015;6:91.

10. Witkowska M, Smolewski P, Robak T. Investigational therapies targeting CD37 for the treatment of B-cell lymphoid malignancies. Expert Opin Investig Drugs. 2018;27(2):171–7.

11. Payandeh Z, Noori E, Khalesi B, Mard-Soltani M, Abdolalizadeh J, Khalili S. Anti-CD37 targeted immunotherapy of B-Cell malignancies. Biotechnol Lett. 2018;40(11-12):1459–66.

12. Link MP, Bindl J, Meeker TC, Carswell C, Doss CA, Warnke RA, et al. A unique antigen on mature B cells defined by a monoclonal antibody. J Immunol. 1986;137(9):3013–8.

13. Schwartz-Albiez R, Dorken B, Hofmann W, Moldenhauer G. The B cell-associated CD37 antigen (gp40-52). Structure and subcellular expression of an extensively glycosylated glycoprotein. J Immunol. 1988;140(3):905–14.

14. Orinska Z, Hagemann PM, Halova I, Draber P. Tetraspanins in the regulation of mast cell function. Med Microbiol Immunol. 2020;209(4):531–43.

15. de Winde CM, Veenbergen S, Young KH, Xu-Monette ZY, Wang XX, Xia Y, et al. Tetraspanin CD37 protects against the development of B cell lymphoma. J Clin Invest. 2016;126(2):653–66.

16. van Spriel AB, de KS, van der Schaaf A, Gartlan KH, Sofi M, Light A, et al. The tetraspanin CD37 orchestrates the alpha(4)beta(1) integrin-Akt signaling axis and supports long-lived plasma cell survival. Sci Signal. 2012;5(250):ra82.

17. Bobrowicz M, Kubacz M, Slusarczyk A, Winiarska M. CD37 in B cell derived tumors—more than just a docking point for monoclonal antibodies. International Journal of Molecular Sciences. 2020;21(24):9531.

18. Lapalombella R, Yeh YY, Wang L, Ramanunni A, Rafiq S, Jha S, et al. Tetraspanin CD37 directly mediates transduction of survival and apoptotic signals. Cancer Cell. 2012;21(5):694–708.

19. Barrena S, Almeida J, Yunta M, Lopez A, Fernandez-Mosteirin N, Giralt M, et al. Aberrant expression of tetraspanin molecules in B-cell chronic lymphoproliferative disorders and its correlation with normal B-cell maturation. Leukemia. 2005;19(8):1376–83.

20. Moore K, Cooper SA, Jones DB. Use of the monoclonal antibody WR17, identifying the CD37 gp40-45 Kd antigen complex, in the diagnosis of B-lymphoid malignancy. J Pathol. 1987;152(1):13–21.

21. Dahle J, Repetto-Llamazares AH, Mollatt CS, Melhus KB, Bruland OS, Kolstad A, et al. Evaluating antigen targeting and anti-tumor activity of a new anti-CD37 radioimmunoconjugate against non-Hodgkin’s lymphoma. Anticancer Res. 2013;33(1):85–95.

22. Stathis A, Flinn IW, Madan S, Maddocks K, Freedman A, Weitman S, et al. Safety, tolerability, and preliminary activity of IMGN529, a CD37-targeted antibody-drug conjugate, in patients with relapsed or refractory B-cell non-Hodgkin lymphoma: a dose-escalation, phase I study. Invest New Drugs. 2018;36(5):869–76.

23. Sawas A, Savage KJ, Perez RP, Advani RH, Melhem-Bertrandt A, Lackey J, et al. A first in human experience of the anti-CD37 antibody-drug conjugate AGS67E in lymphoid malignancies. Journal of Clinical Oncology. 2016;34(15_suppl):7549-.

24. Levy MY, Jagadeesh D, Grudeva-Popova Z, Trněný M, Jurczak W, Pylypenko H, et al. Safety and Efficacy of CD37-Targeting Naratuximab Emtansine PLUS Rituximab in Diffuse Large B-Cell Lymphoma and Other NON-Hodgkin’S B-Cell Lymphomas-a Phase 2 Study. Blood. 2021;138:526.

25. Repetto-Llamazares AH, Larsen RH, Patzke S, Fleten KG, Didierlaurent D, Pichard A, et al. Targeted Cancer Therapy with a Novel Anti-CD37 Beta-Particle Emitting Radioimmunoconjugate for Treatment of Non-Hodgkin Lymphoma. PLoS One. 2015;10(6):e0128816.

26. Kolstad A, Illidge T, Bolstad N, Spetalen S, Madsbu U, Stokke C, et al. Phase 1/2a study of 177Lu-lilotomab satetraxetan in relapsed/refractory indolent non-Hodgkin lymphoma. Blood Advances. 2020;4(17):4091–101.

27. Zhao X, Lapalombella R, Joshi T, Cheney C, Gowda A, Hayden-Ledbetter MS, et al. Targeting CD37-positive lymphoid malignancies with a novel engineered small modular immunopharmaceutical. Blood. 2007;110(7):2569–77.

28. Heider KH, Kiefer K, Zenz T, Volden M, Stilgenbauer S, Ostermann E, et al. A novel Fc-engineered monoclonal antibody to CD37 with enhanced ADCC and high proapoptotic activity for treatment of B-cell malignancies. Blood. 2011;118(15):4159–68.

29. Oostindie SC, van der Horst HJ, Kil LP, Strumane K, Overdijk MB, van den Brink EN, et al. DuoHexaBody-CD37((R)), a novel biparatopic CD37 antibody with enhanced Fc-mediated hexamerization as a potential therapy for B-cell malignancies. Blood Cancer J. 2020;10(3):30.

30. Scarfo I, Ormhoj M, Frigault MJ, Castano AP, Lorrey S, Bouffard AA, et al. Anti-CD37 chimeric antigen receptor T cells are active against B- and T-cell lymphomas. Blood. 2018;132(14):1495–506.

31. Singh V, Gupta D, Almasan A. Development of Novel Anti-Cd20 Monoclonal Antibodies and Modulation in Cd20 Levels on Cell Surface: Looking to Improve Immunotherapy Response. J Cancer Sci Ther. 2015;7(11):347–58.

32. von Horsten HH, Ogorek C, Blanchard V, Demmler C, Giese C, Winkler K, et al. Production of non-fucosylated antibodies by co-expression of heterologous GDP-6-deoxy-D-lyxo-4-hexulose reductase. Glycobiology. 2010;20(12):1607–18.

33. Labrijn AF, Meesters JI, Priem P, de Jong RN, van den Bremer ET, van Kampen MD, et al. Controlled Fab-arm exchange for the generation of stable bispecific IgG1. Nat Protoc. 2014;9(10):2450–63.

34. Skånland SS. Phospho flow cytometry with fluorescent cell barcoding for single cell signaling analysis and biomarker discovery. JoVE (Journal of Visualized Experiments). 2018(140):e58386.

35. Andreatta M, Karosiene E, Rasmussen M, Stryhn A, Buus S, Nielsen M. Accurate pan-specific prediction of peptide-MHC class II binding affinity with improved binding core identification. Immunogenetics. 2015;67(11-12):641–50.

36. Schaffer AA, Aravind L, Madden TL, Shavirin S, Spouge JL, Wolf YI, et al. Improving the accuracy of PSI-BLAST protein database searches with composition-based statistics and other refinements. Nucleic Acids Res. 2001;29(14):2994–3005.

37. Motulsky H, Neubig R. Analyzing radioligand binding data. Curr Protoc Protein Sci. 2001;Appendix 3:Appendix.

38. Repetto-Llamazares AHV, Malenge MM, O’Shea A, Eiriksdottir B, Stokke T, Larsen RH, et al. Combination of (177) Lu-lilotomab with rituximab significantly improves the therapeutic outcome in preclinical models of non-Hodgkin’s lymphoma. Eur J Haematol. 2018;101(4):522–31.

39. Motulsky H, Neubig R. Analyzing radioligand binding data. Curr Protoc Neurosci. 2002;Chapter 7:Unit 7 5.

40. Maaland AF, Heyerdahl H, O’Shea A, Eiriksdottir B, Pascal V, Andersen JT, et al. Targeting B-cell malignancies with the beta-emitting anti-CD37 radioimmunoconjugate (177)Lu-NNV003. Eur J Nucl Med Mol Imaging. 2019;46(11):2311–21.

41. Grevys A, Bern M, Foss S, Bratlie DB, Moen A, Gunnarsen KS, et al. Fc Engineering of Human IgG1 for Altered Binding to the Neonatal Fc Receptor Affects Fc Effector Functions. J Immunol. 2015;194(11):5497–508.

42. Foss S, Grevys A, Sand KMK, Bern M, Blundell P, Michaelsen TE, et al. Enhanced FcRn dependent transepithelial delivery of IgG by Fc-engineering and polymerization. J Control Release. 2016;223:42–52.

43. Grevys A, Nilsen J, Sand KMK, Daba MB, Oynebraten I, Bern M, et al. A human endothelial cell-based recycling assay for screening of FcRn targeted molecules. Nat Commun. 2018;9(1):621.

44. Roopenian DC, Christianson GJ, Proetzel G, Sproule TJ. Human FcRn Transgenic Mice for Pharmacokinetic Evaluation of Therapeutic Antibodies. Methods Mol Biol. 2016;1438:103–14.

45. Hosseini I, Gajjala A, Bumbaca Yadav D, Sukumaran S, Ramanujan S, Paxson R, et al. gPKPDSim: a SimBiology((R))-based GUI application for PKPD modeling in drug development. J Pharmacokinet Pharmacodyn. 2018;45(2):259–75.

46. Hamze M, Meunier S, Karle A, Gdoura A, Goudet A, Szely N, et al. Characterization of CD4 T Cell Epitopes of Infliximab and Rituximab Identified from Healthy Donors. Front Immunol. 2017;8:500.

47. Kim SY, Theunissen JW, Balibalos J, Liao-Chan S, Babcock MC, Wong T, et al. A novel antibody-drug conjugate targeting SAIL for the treatment of hematologic malignancies. Blood Cancer J. 2015;5:e316.

48. Golay J, Taylor RP. The Role of Complement in the Mechanism of Action of Therapeutic Anti Cancer mAbs. Antibodies (Basel). 2020;9(4).

49. Overdijk MB, Verploegen S, Ortiz BA, Vink T, Leusen JH, Bleeker WK, et al. Crosstalk between human IgG isotypes and murine effector cells. J Immunol. 2012;189(7):3430–8.

50. Herting F, Friess T, Bader S, Muth G, Holzlwimmer G, Rieder N, et al. Enhanced anti-tumor activity of the glycoengineered type II CD20 antibody obinutuzumab (GA101) in combination with chemotherapy in xenograft models of human lymphoma. Leuk Lymphoma. 2014;55(9):2151–5160.

51. Saxena A, Wu D. Advances in Therapeutic Fc Engineering - Modulation of IgG-Associated Effector Functions and Serum Half-life. Front Immunol. 2016;7:580.

52. Wright A, Morrison SL. Effect of altered CH2-associated carbohydrate structure on the functional properties and in vivo fate of chimeric mouse-human immunoglobulin G1. J Exp Med. 1994;180(3):1087–96.

53. Kanda Y, Yamada T, Mori K, Okazaki A, Inoue M, Kitajima-Miyama K, et al. Comparison of biological activity among nonfucosylated therapeutic IgG1 antibodies with three different N-linked Fc oligosaccharides: the high-mannose, hybrid, and complex types. Glycobiology. 2007;17(1):104–18.

54. Bas M, Terrier A, Jacque E, Dehenne A, Pochet-Beghin V, Beghin C, et al. Fc Sialylation Prolongs Serum Half-Life of Therapeutic Antibodies. J Immunol. 2019;202(5):1582–94.

55. Morell AG, Gregoriadis G, Scheinberg IH, Hickman J, Ashwell G. The role of sialic acid in determining the survival of glycoproteins in the circulation. J Biol Chem. 1971;246(5):1461–7.

56. Xu Y, Chen G. Mast cell and autoimmune diseases. Mediators Inflamm. 2015;2015:246126.

57. de Gruijter NM, Jebson B, Rosser EC. Cytokine production by human B cells: role in health and autoimmune disease. Clin Exp Immunol. 2022.

