## Supplementary tables and figures for "NNV024, a novel humanized anti-CD37 antibody with enhanced ADCC and prolonged plasma half-life in human FcRn transgenic mice for treatment of NHL"

### Supplementary material

**Supplementary figure 1**

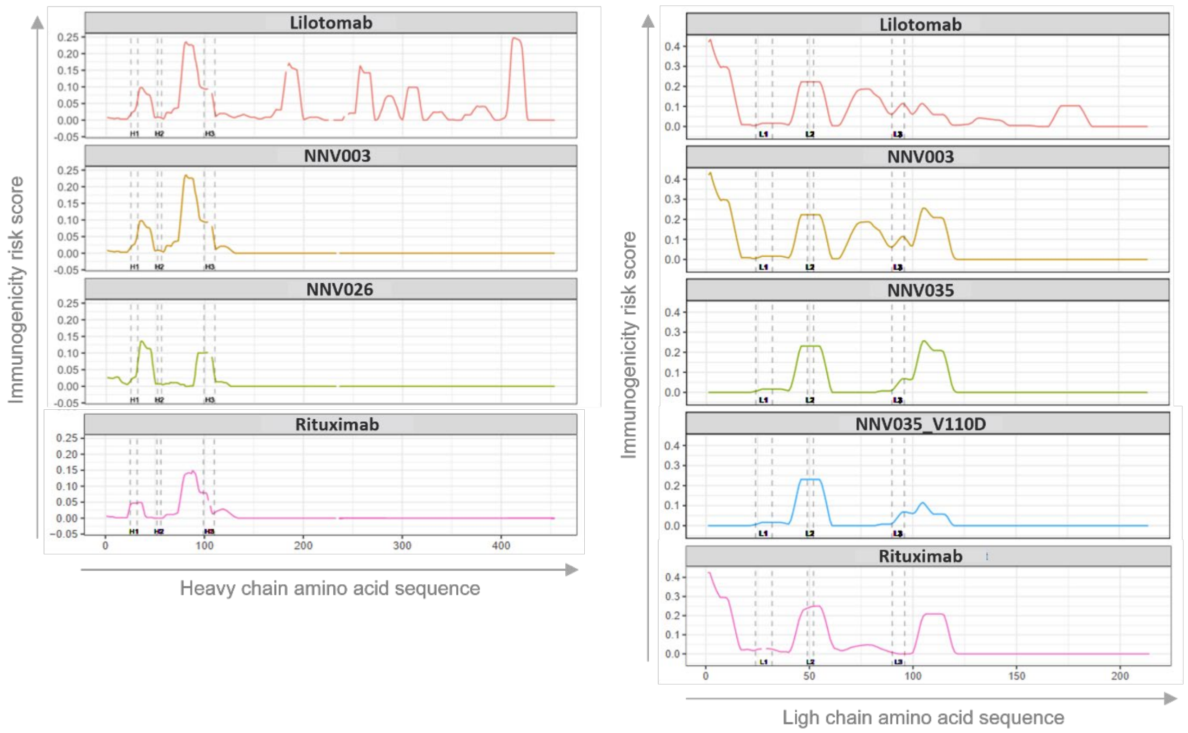


**Supplementary figure 1**. Self-adjusted position-specific immunogenicity risk profiles

Illustration of the self-adjusted position-specific immunogenicity risk scores calculated for the world population for the heavy (Left panel) and light (Right panel) chains of lilotomab, NNV003, humanized derivatives hereof. Rituximab is added for benchmarking purposes. Dotted lines highlight the position of the complementary determining regions (CDR, H1-3 for HC and L1-3 for the LC from left to right) determined using a trained hidden Markov model.

**Supplementary Table 1**. Binding to human FcγRs and FcRn at pH 6

| MAb variant | EC50 (ng/mL) | | | | | | |
|---|---|---|---|---|---|---|---|
| | FcγRIIIa | | FcγRIIa | | FcγRIIb | FcγRIIIb | FcγRI |
| | -F158 | -V158 | -H131 | -R131 | | | |
| NNV025 | 534.7 | 987.7 | 893.5 | 803.3 | 779.4 | 1323 | 660.1 |
| NNV023 | 460.8 | 860.7 | 761.9 | 706.2 | 649.6 | 1370 | 646.4 |
| NNV024 | 104.1 | 332.4 | 766.6 | 689.7 | 674.3 | 484.3 | 588.2 |
| obinutuzumab | 296.5 | 1150.0 | 2298.0 | 2113.0 | 2801.0 | 2456 | 1981.0 |
| DuoHexaBody-CD37 | 2150 | 4166.0 | 4392.0 | 4095.0 | 3439.0 | n.d.* | 3191.0 |

*n.d. = not determined

#### Supplementary figure 2

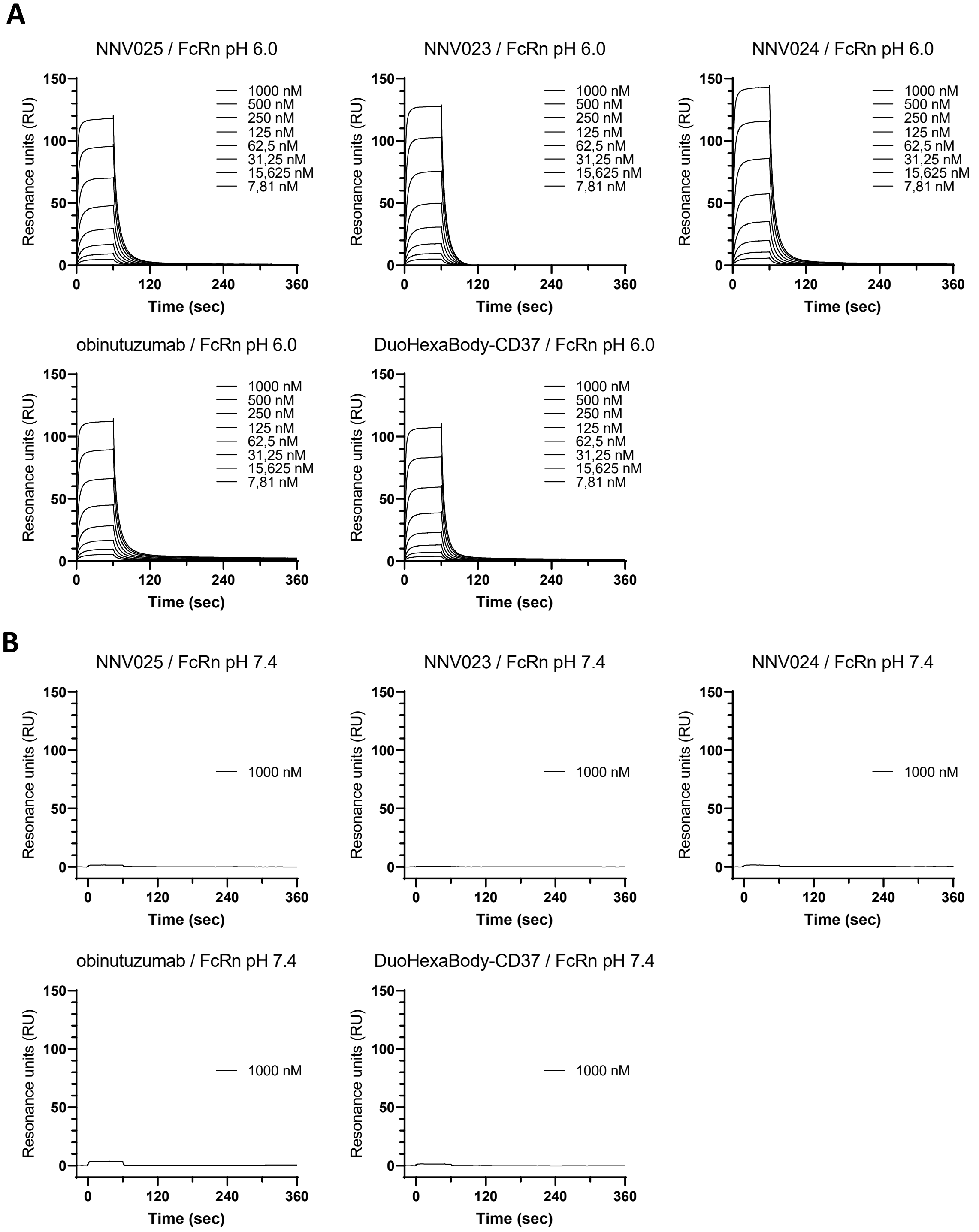


**Supplementary figure 2.** SPR binding kinetics to hFcRn. The sensorgrams show binding of FcRn-His6x to immobilized test antibodies at pH 6.0 (1000 – 7.91 nM, panel A) and pH 7.4 (1000 nM only, panel B).

**Supplementary figure 3**

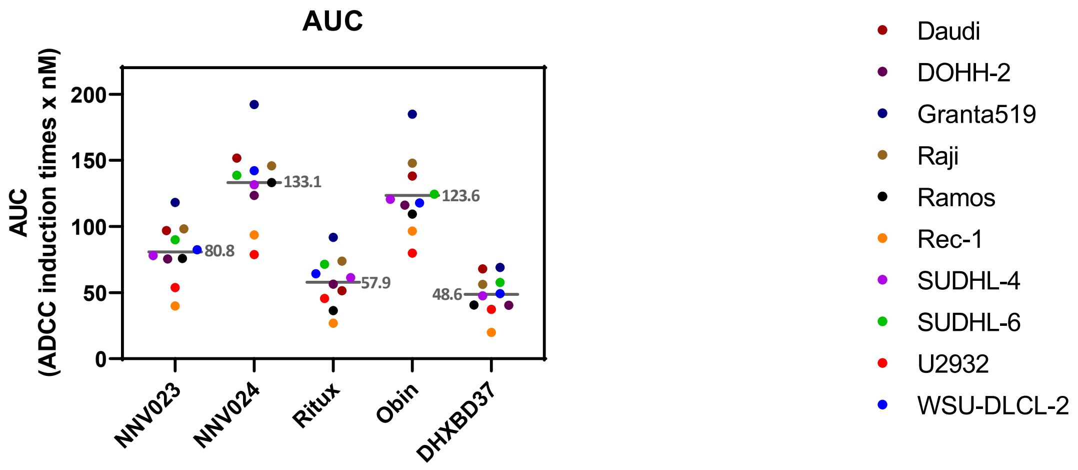


**Supplementary figure 3**. Area Under the Curve (AUC) for the ADCC concentration response curves. The grey dash in each group of scatter represents the mean of the group, the numeric value of the mean is displayed next to it. The results for REH cell line are not shown since the cell line does not express CD20 and CD37 to a sufficient level to generate a concentration-response curve. Abbreviations: Ritux – rituximab; Obin – obinutuzumab.

**Supplementary figure 4**

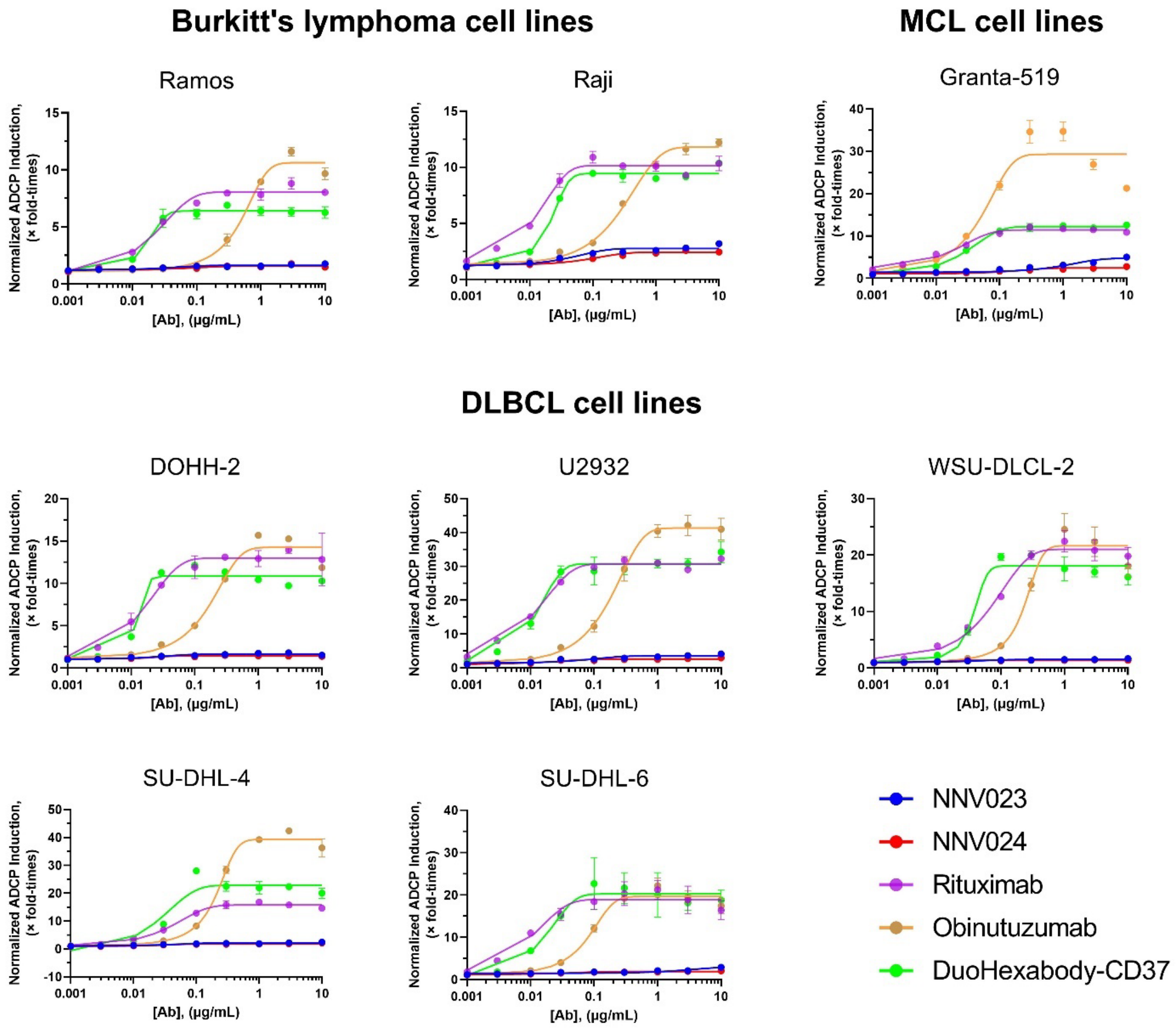


**Supplementary Figure 4. Normalized ADCP induced by test antibodies incubated with FcγRIIa-131R expressing effector cells in a panel of NHL cell lines.** Relative ADCP induction was assessed in a panel of ten NHL cell lines using the ADCP FcγRIIa-131R reporter assay (Promega). Bioluminescent signal is obtained through FcγRIIa/NFAT-associated luciferase activation in FcγRIIa-131R expressing effector cells. The ADCP reporter assay response is plotted in function of the antibody concentration. The result is the mean of two technical replicates normalized to the untreated within-the-plate control (untreated Target + Effector cells). The error bars are SD. The spline lines are the fit of the data to a sigmoidal 4PL curve. Abbreviations: DLBCL – Diffused Large B-Cell Lymphoma; MCL – Mantle Cell Lymphoma.

**Supplementary figure 5**

#### Burkitt Lymphoma's cell lines

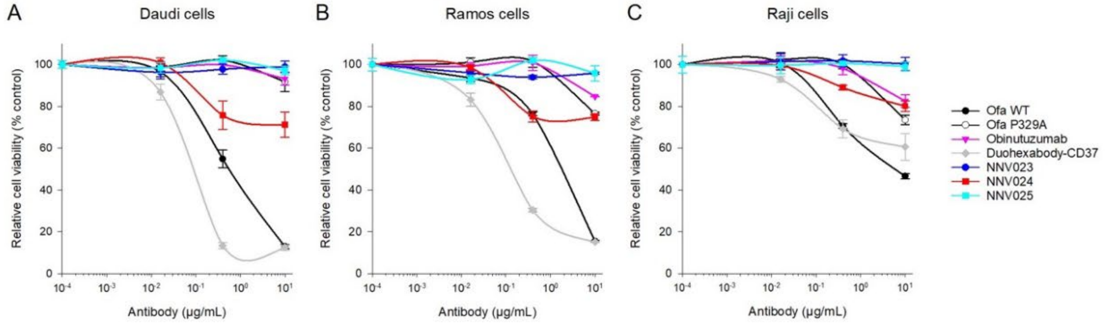


#### DLBCL cell lines

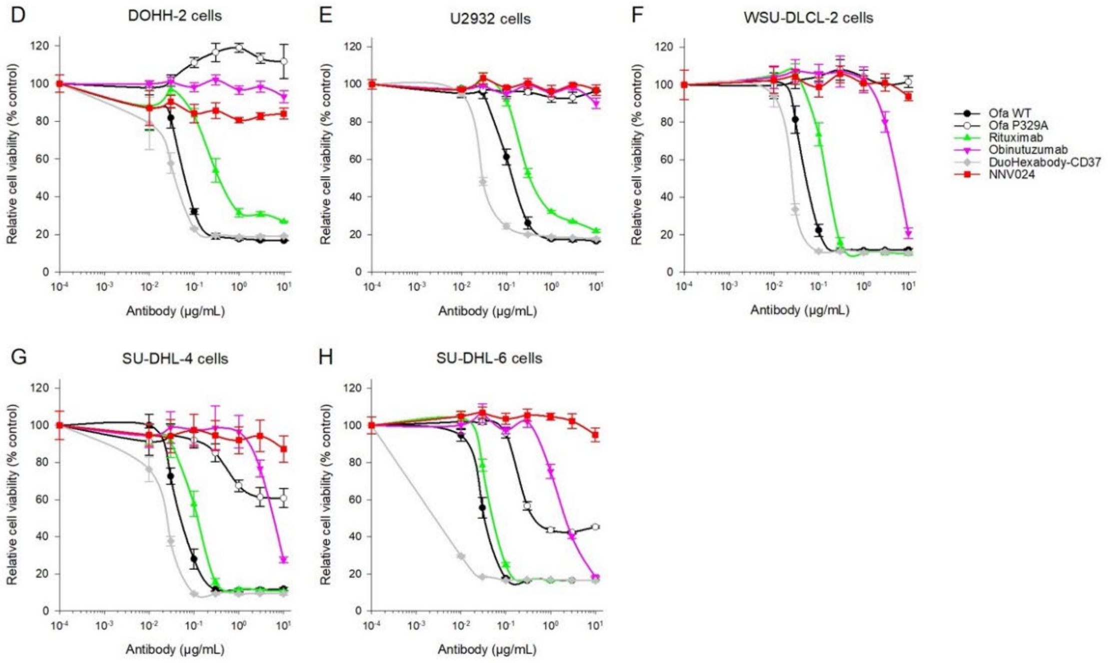


#### MCL cell lines

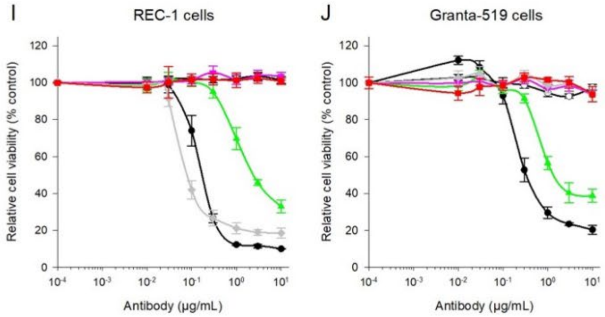

**Supplementary Figure 5** Assessment of antibody-induced CDC on a panel of NHL cell lines. Relative cell viability upon treatment with different concentrations of test antibodies (serial dilutions 1:3, concentration range 0.0001-10 µg/mL) in a panel of 3 Burkitt's Lymphoma, 5 DLBCL and 2 MCL cell lines. The cell viability was quantified as Alamar Blue-induced fluorescent signal and normalized on untreated control cell signal. The relative cell viability values are expressed as percentage of control (y axis) and are plotted in function of the Ab concentration (x axis) to generate dose-response curves. The results represented are the mean of three technical replicates and the error bars are SD. The spline lines are the fit of the data to a sigmoidal 4PL curve. Abbreviations: DLBCL – Diffused Large B-Cell Lymphoma; MCL – Mantle Cell Lymphoma; Ritux – rituximab; Obin – obinutuzumab; DHXBD37 – DuoHexaBody-CD37; Ofa WT – ofatumumab wild-type; Ofa P329A – ofatumumab P329A mutant.

**Supplementary Table 2**. Plasma half-life assessed by percentage of mAb remaining in plasma

| Groups | Plasma half-life (days) |
|---|---|
| NNV025 | 7.4 ± 0.8 |
| NNV023 | 12.0 ± 2.4* |
| NNV024 | 8.9 ± 0.8* |
| obinutuzumab | 4.4 ± 1.3 |
| DuoHexabody-CD37 | 4.1 ± 1.2 |

*Significantly different from obinutuzumab and DuoHexaBody-CD37 (t-test, $p < 0.01$)

**Supplementary Table 3**. **Median survival** of CB17-SCID mice with i.v. injected Daudi cells treated with 1, 3 or 6 injections of 100 µg NNV024 or obinutuzumab or 100 µl NaCl (N = 10 mice per group).

| Treatment group | Median Survival (days from inoculation) |
|---|---|
| NNV024 – 100 µg x 1 | 85 |
| NNV024 – 100 µg x 3 | 79 |
| NNV024 – 100 µg x 6 | 80 |
| Obinutuzumab – 100 µg x 1 | 88 |
| Obinutuzumab – 100 µg x 3 | 90 |
| Obinutuzumab – 100 µg x 6 | 77 |
| 0.9% NaCl x 3 | 24 |

#### Supplementary Figure 6

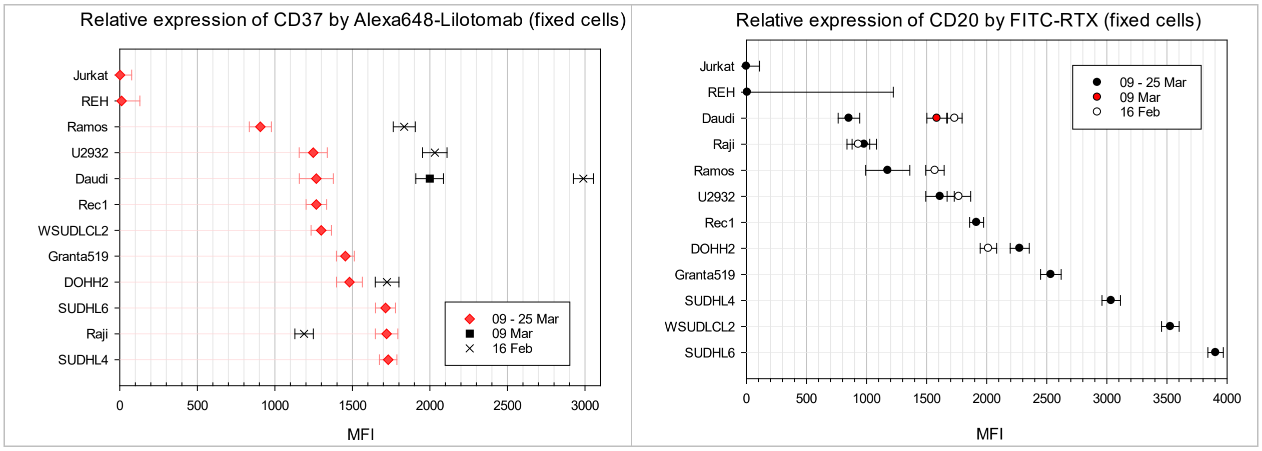


**Supplementary Figure 6**. Relative expression levels of CD37 in NHL lines detected with Alexa648-labelled lilotomab (A) and expression of CD20 in NHL lines detected with FITC-labelled rituximab. A drift in the expression level depending on the passage number have been noticed. All NHL cells lines used in the study were in the culture for no more than 30 passages in total (less than 10 weeks).

#### Supplementary Figure 7

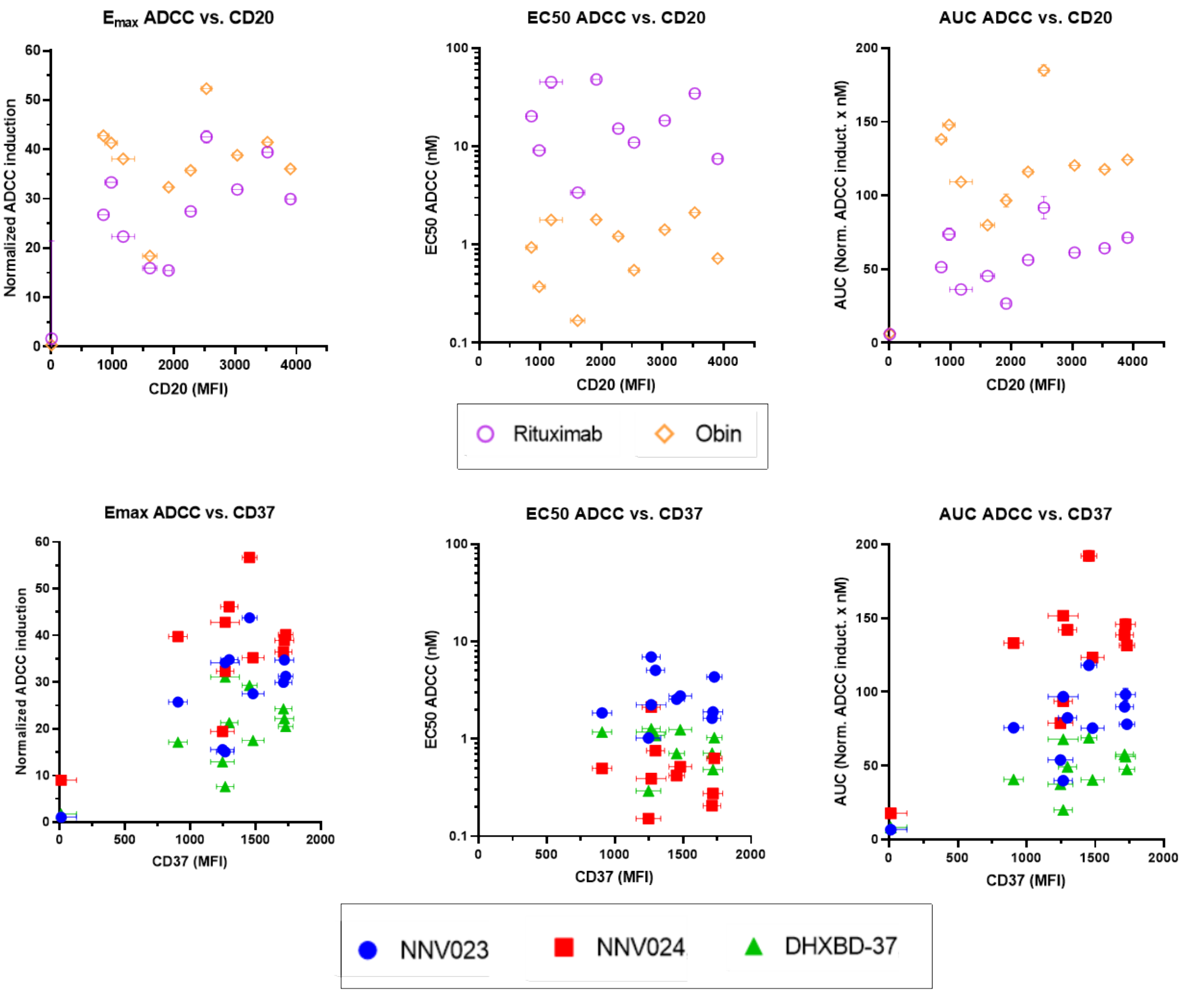


**Supplementary Figure 7**. ADCC parameters (Emax, $EC_{50}$, AUC) of the *in vitro* tested antibodies do not correlate with the amount of CD37 or CD20 expressed on the NHL cells used in the study. The symbols are the mean fluorescence intensity, the bars are the standard deviation of MFI.

**Supplementary Figure 8**

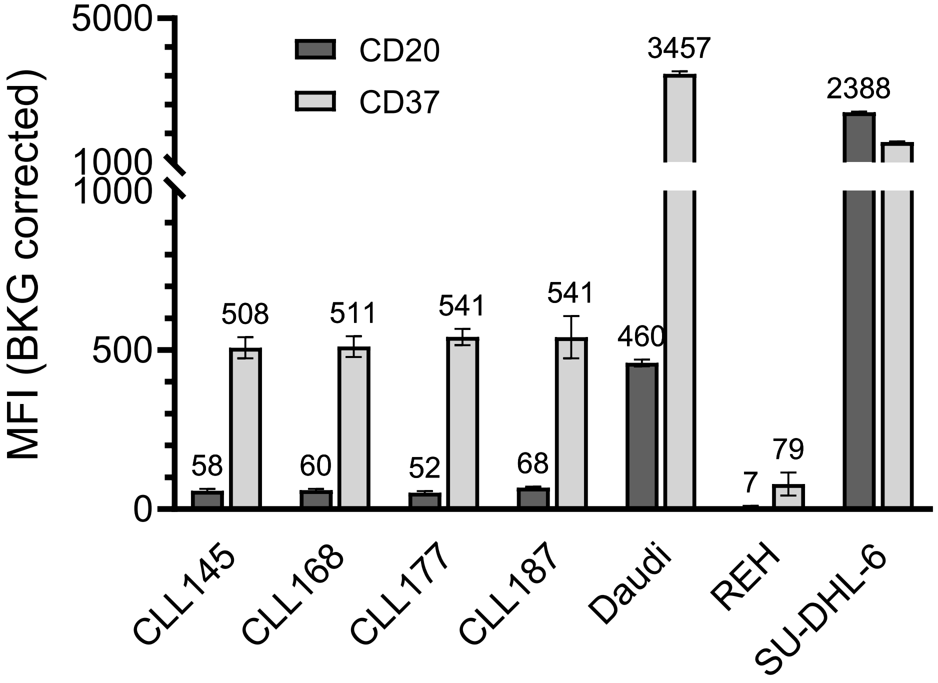


**Supplementary Figure 8**. Relative expression levels of CD37 and CD20 in the patient derived CLL samples and NHL cell lines used as expression level controls. Alexa648-labelled l lilotomab was used for detecting CD37, while the FITC-labelled rituximab was used for detecting CD20.

**Supplementary Figure 9**

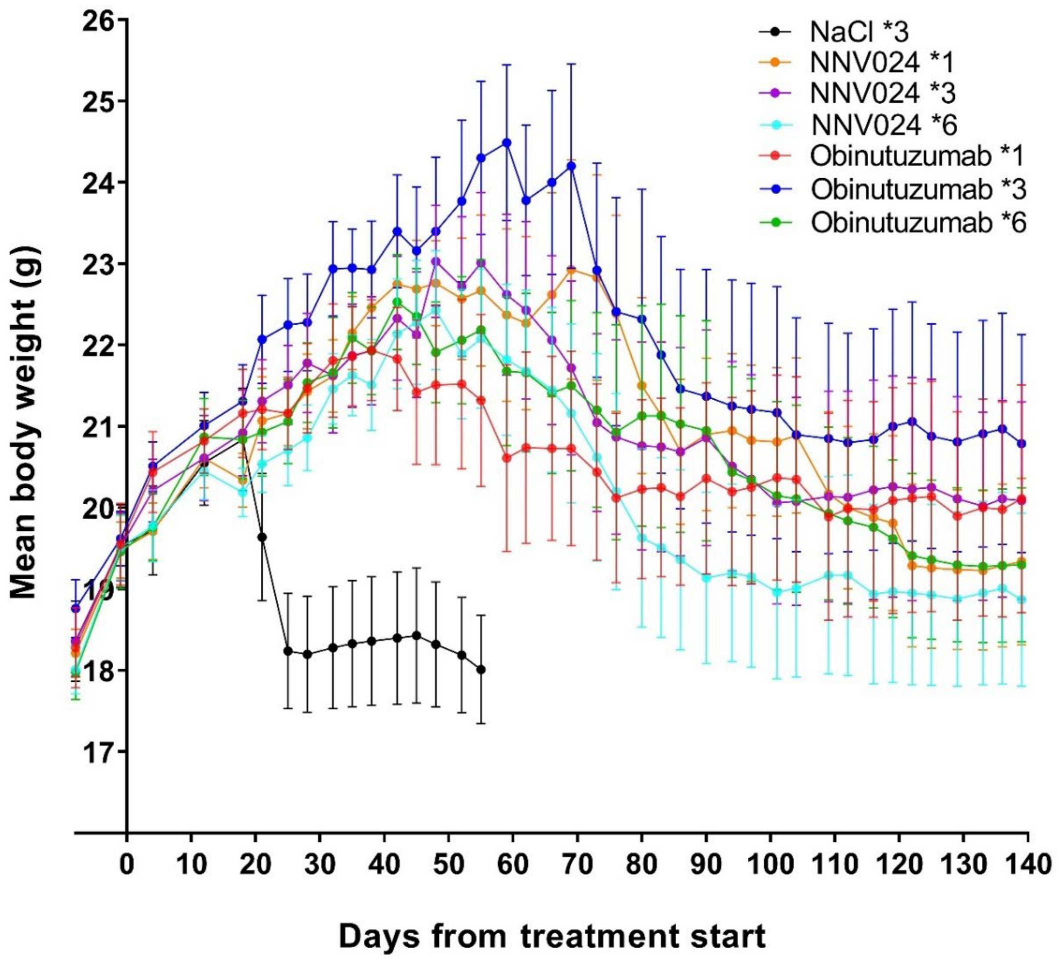


**Supplementary Figure 9.** Mean body weight of CB17-SCID mice with i.v. injected Daudi cells treated with 1, 3 or 6 injections of 100 μg NNV024 or obinutuzumab or 100 μl NaCl (N = 10 mice per group). Error bars indicate standard error of the mean.

**Supplementary Table 4. Median survival** of CB17-Scid mice with i.v. injected Daudi cells (1x$10^7$ cells and treated with 10 or 50 µg/mouse of NNV024 or obinutuzumab or 100 ml NaCl (N=10 mice per group).

| **Treatment group** | **Median Survival (days from inoculation)*** | **p-value summary vs. NNV024 – 50 µg** | **% animals surviving 118 days after inoculation without signs of tumors** |
|---|---|---|---|
| NNV024 – 50 µg | 84 | - | 30 |
| NNV024 – 10 µg | 70.5 | n.s. | 20 |
| Obinutuzumab – 50 µg | 62 | 0.0157 | 0 |
| Obinutuzumab -10 µg | 53 | 0.0119 | 10 |
| 0.9% NaCl | 23.5 | < 0.0001 | 0 |

*All treatment groups had significantly better survival than the control group (0.9% NaCl), $p < 0.0001$, Mantel-Cox Test. The pairwise comparison of the test groups was performed according to the Holm-Sidak method (α=0.05).

**Supplementary Figure 10**

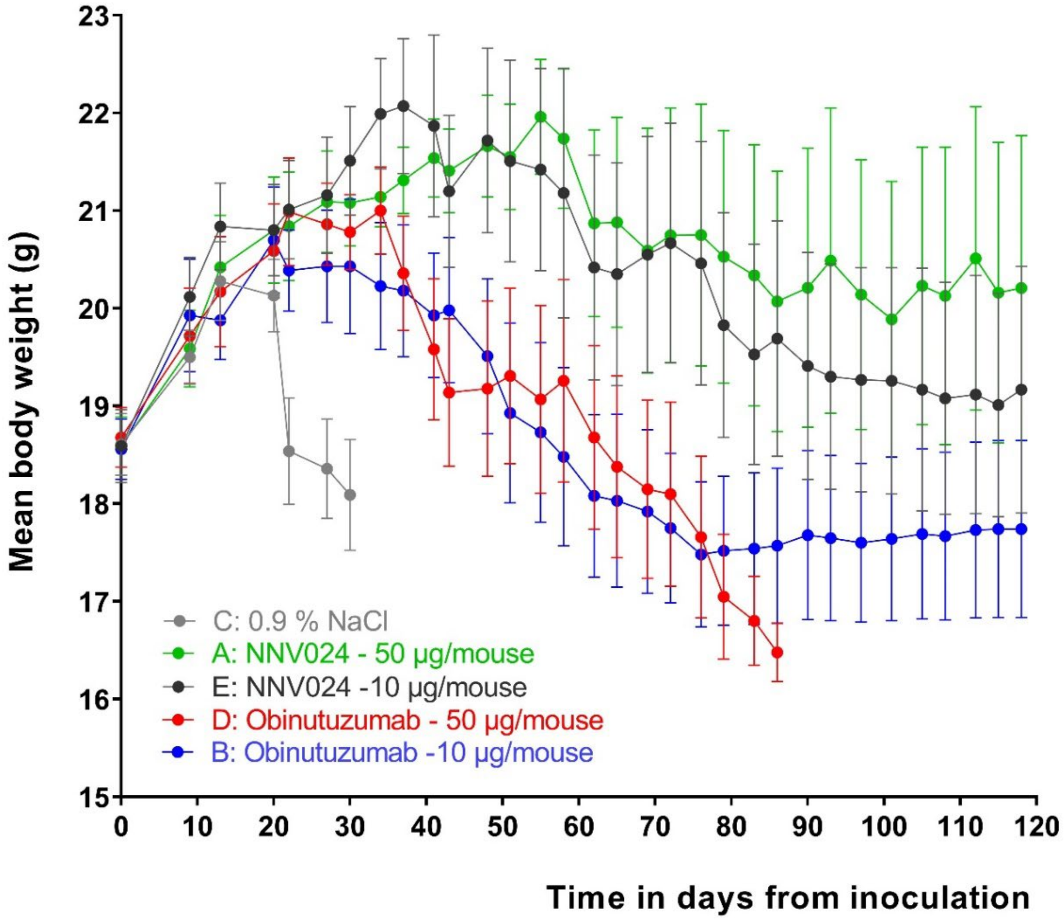


**Supplementary Figure 10**. Mean body weight of CB17-Scid mice with i.v. injected Daudi cells and treated with 10 or 50 μg/mouse of NNV024 or obinutuzumab or 100 ml NaCl (N=10 mice per group). Error bars indicate standard error of the mean.

**Supplementary Figure 11**

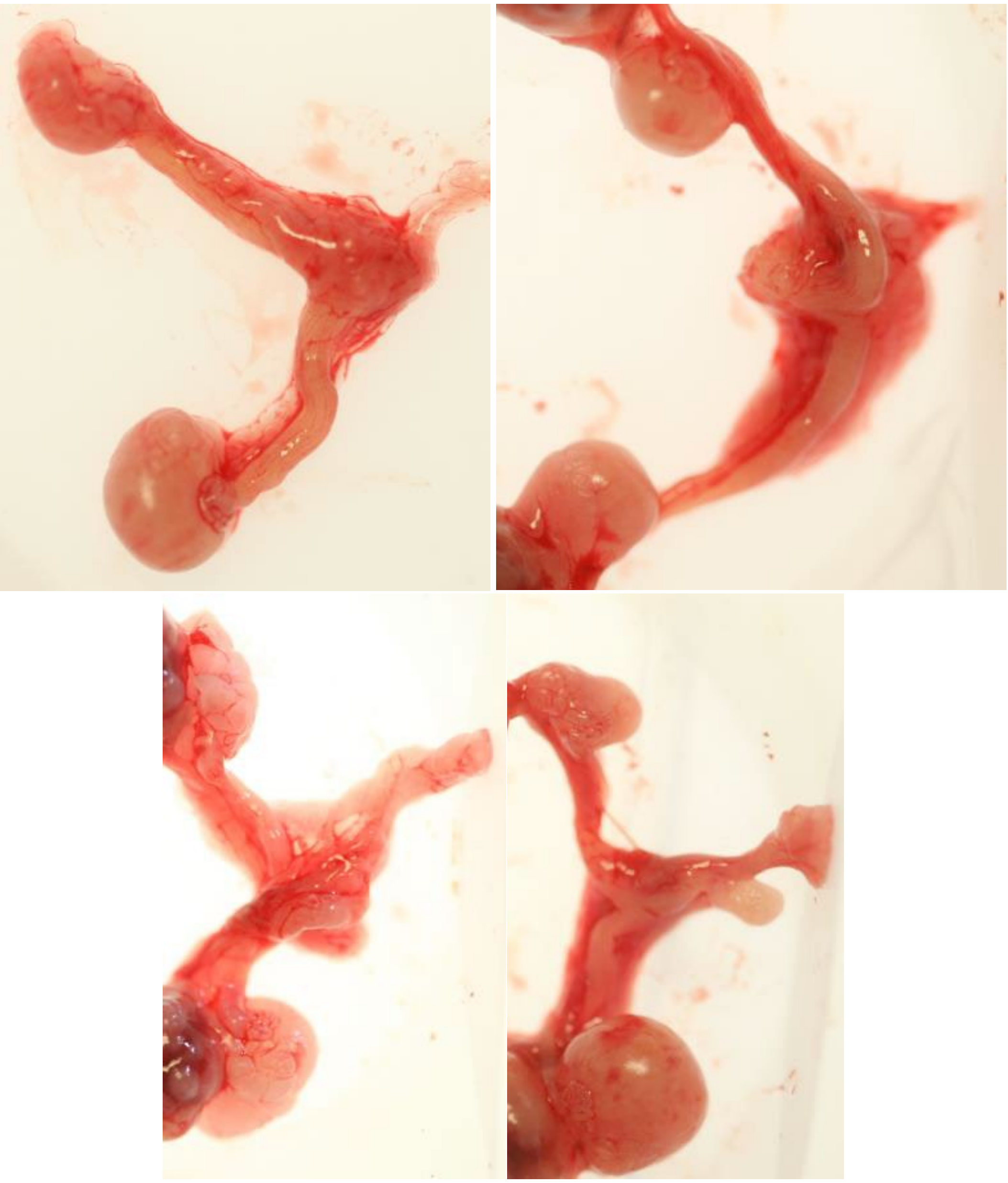

**Supplementary Figure 11.** Pictures of Daudi-induced ovarian tumors from 4 mice in the study.

#### Supplementary Figure 12

**A. Sparse amount of small nodular tumors (< 10)**

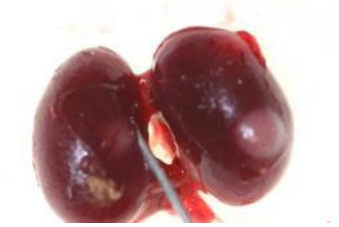

K1

**K1:** Less than 10 small nodular tumors on the surface of the kidney.

**B. Multiple (>10) small nodular tumors, giving the kidneys a grape like appearance.**

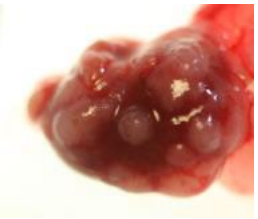

K2

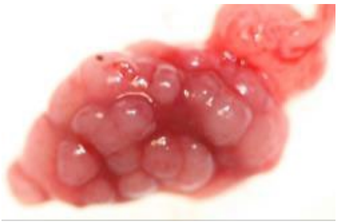

K3

**K2:** Multiple (>10) small nodular tumors with some normal kidney tissue visible.

**K3:** Multiple (>10) small nodular tumors with sparse normal kidney tissue visible.

**C. Multiple (>10) large nodular tumors**

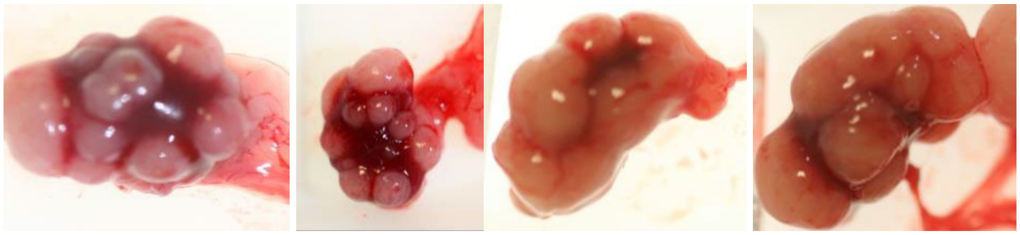

K4 K5

**K4:** Numerus large nodular tumors with clear boundaries and some normal kidney tissue visible.

**K5:** Numerus large nodular tumors with unclear boundaries (the noduli have fused together) and sparse normal kidney tissue visible.

**D. Deformed large kidney tumors**

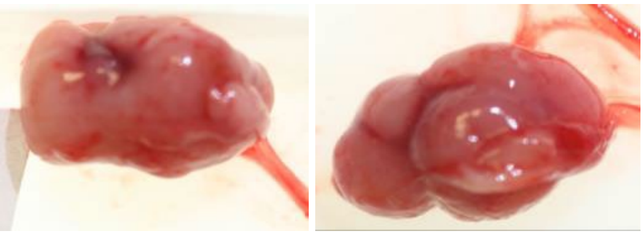

**K6**

**K6:** Large Kidney tumors, noduli have fused together to form a large tumor mass.

**E. Monstrous kidney tumors**

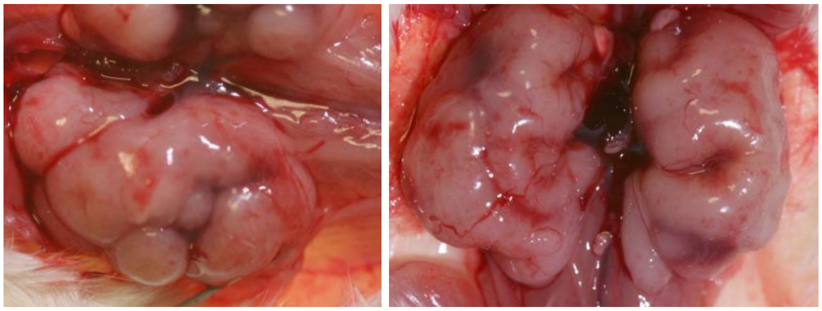

K7

**K7:** Kidney that have transformed into huge tumor mass.

**Supplementary Figure 12.** Pictures of Daudi-induced kidney tumors.
